# Single-B cell analysis correlates high-lactate secretion with stress and increased apoptosis

**DOI:** 10.1101/2023.09.01.555863

**Authors:** Olivia T.M. Bucheli, Daniela Rodrigues, Kevin Portmann, Aline Linder, Marina Thoma, Cornelia Halin, Klaus Eyer

## Abstract

While cellular metabolism was proposed to be a driving factor of the activation and differentiation of B cells and the function of the resulting antibody-secreting cells (ASCs), the study of correlations between cellular metabolism and functionalities has been difficult due to the absence of technologies enabling the parallel measurement. Herein, we performed single-cell transcriptomics and introduced a direct concurrent functional and metabolic flux quantitation of individual murine B cells. Our transcriptomic data identified lactate metabolism as dynamic in ASCs, but antibody secretion did not correlate with lactate secretion rates (LSRs). Instead, our study of all splenic B cells during an immune response linked increased lactate metabolism with acidic intracellular pH and the upregulation of apoptosis. T cell-dependent responses increased LSRs, and added TLR4 agonists affected the magnitude and boosted LSR^high^ B cells *in vivo*, while resulting in only a few immunoglobulin-G secreting cells (IgG-SCs). Therefore, our observations indicated that LSR^high^ cells were not differentiating into IgG-SCs, and were rather removed due to apoptosis.

## Introduction

Vaccine-induced protection by humoral immunity involves generating and maintaining immunoglobulin G-secreting cells (IgG-SCs) as a final goal for long-lasting protection. IgG-SCs arise from antigen-activated B cells within secondary lymphoid organs such as the spleen. This process is initiated by B cell receptor (BCR) binding to the antigen with potential additional co-stimulatory signaling by CD40, interleukin-4, and Toll-like receptors (TLRs) [1-4]. Reactive oxygen species (ROS) production upon BCR and co-receptor activation was shown to play a crucial role by increasing and prolonging the signal [5]. B cells undergo clonal expansion upon activation, demanding a rapid adjustment of the metabolic program to support cellular growth and proliferation [6], upregulating glycolysis and oxidative phosphorylation (OXPHOS) [7-14]. A fraction of cells rapidly starts to produce antibodies in a short-lasting extrafollicular response [15-18], whereas other B cells give rise to germinal center (GC) B cells [19, 20], which undergo extensive proliferation, affinity maturation, and selection [20, 21]. Current literature suggests that GC B cell’s metabolism is highly heterogeneous, dynamic and that metabolites play a variety of roles beyond bioenergetics [13, 14, 22-30]. In addition, hypoxia was observed in the GCs [22, 27], again underlying a changing environment and the need for adaptation from the GC B cell’s side.

Ultimately, the GC response outputs high-affine effector cells, either memory B cells or long-lived antibody-secreting cells (ASCs). FAO was shown to be the main energy source of long-lived IgG-SCs [31], while amino acids and glucose were mainly used to produce antibodies. Stress responses in ASCs increased autophagy and the antioxidant defense system to cope with potentially lethal stresses due to ROS and the misfolding of proteins [32, 33]. Glucose also contributed to overcoming stressful metabolic situations in long-lived ASCs [31]. Additionally, the secreted antibody isotype seemed to play a factor in metabolism, as a preference for glycolysis for IgA-secreting cells was described in literature [34], although these differences might also be niche-dependent. Most long-lived IgG-SCs migrate to the bone marrow, where they reside in survival niches [35, 36], which are known for being dynamic environments with varying nutritional conditions [37]. Therefore, there are various complex interplays between cellular metabolism, formation, function, and survival of IgG-SCs.

Various technologies for studying metabolic reprogramming, metabolic status, and metabolic fluxes are available [38]. Transcriptomic analyses on the single-cell level have contributed enormously to our current understanding of metabolic regulation and have been particularly useful for identifying differences in gene expression between immune cell populations [9, 10, 22, 29-31, 39-41]. However, well-described discrepancies between mRNA level and protein activity can occur due to post-transcriptional, translational, and allosteric regulations [42-44], and consequently, transcript analysis only assesses metabolism as a fold-change in expression. Alternatively, cellular energy levels can be investigated by assessing protein synthesis levels using a method called SCENITH [45]. Thereby, the contribution of different metabolic pathways can be assessed using distinct inhibitors [45]. However, this method is an indirect measurement and, accordingly, only indicates active pathways. Nevertheless, direct measurements of metabolic pathways can be performed by making use of the uptake of fluorescent derivatives of nutrients like glucose or by measuring oxygen consumption, extracellular acidification rates, or lactate secretion [7, 30, 31, 46, 47]. However, these technologies study cells in bulk or cannot easily be combined with functional studies such as antibody secretion. Indeed, the cellular outputs of IgG-SCs were very heterogenous and varied between 10-3’000 IgG/s [48], differences that might also alter the stress status and metabolic requirements of the individual cells. Therefore, conducting a parallel analysis of metabolic and functional characteristics at the single-cell level could offer valuable insights into complex interplays between cellular metabolism, functionality, and survival of IgG-SCs and differentiating B cells.

In this study, we combined transcriptomics with a parallel direct and quantitative assessment of metabolic fluxes and functionalities of individual B cells. We applied the methods to studying B cells and IgG-SCs at various stages, functions, and metabolic requirements throughout a murine secondary immune response. Due to the ambivalent role of glucose and potential hypoxia in the GCs, we focused on droplet-based measurements of lactate secretion rates (LSRs). For functionalities, we aimed to study IgG secretion rates and affinities, and by advancing the droplet-based microfluidics system ‘DropMap’ [48, 49] to study potential correlations on the single-B cell level. While far from a complete functional assessment of all metabolic fluxes present in individual B cells, the integrated assay provided a starting point to characterize a key metabolic flux present in B cells at various stages on the single-cell level. Our study identified lactate metabolism and LSRs as a new potential influencer of B cell functionality and survival, showing correlations between increased LSRs and low intracellular pH and expression levels of lactate dehydrogenase and apoptotic genes.

## Results

### Metabolic pathways in IgG-ECs are distinct, and a decrease in lactate metabolism is observed in the early immune response

Firstly, we characterized the induced immune response functionally and, in addition, transcriptionally on the single-cell level. We immunized female BALB/c mice with tetanus toxoid (TT) in alum with added Monophosphoryl Lipid A from *Salmonella minnesota* R595 (MPLA), called adjuvant system 04 (AS04). The mice were immunized twice, four weeks apart, and the spleen and bone marrow B cells were analyzed on days 0 (i.e., on the day of secondary immunization, but without application of the immunization), 3, 7 and 14 after secondary immunization (Figure 1A). Using an adapted version of our previously published droplet-based system, IgG-SCs were identified, and their IgG secretion rate and affinity assessed [48, 49]. For this purpose, a sandwich immunoassay was used whereby a fluorescent IgG probe and fluorescently labeled TT relocated onto functionalized paramagnetic nanoparticles in a concentration- and affinity-dependent manner, respectively (Figure 1B). Secretion rates were calculated by monitoring concentration changes over time within the assay’s quantitative range (3-375 IgG/s, SIFigure 1), and cells with a secretion rate ≥3 IgG/s were defined as IgG-SCs. The strength of the interaction between the secreted IgG and the antigen was determined by the slope when plotting the relocation signal of the IgG probe against the relocation signal of the fluorescently labeled TT (SIFigure 1), with increasing slopes indicating higher affinities. The frequency of IgG-SCs remained surprisingly constant and low at the measured days in the spleen but increased and remained stable in the bone marrow after day 7 (Figure 1C). In both compartments, the median IgG secretion rate diminished over the immune response (Figure 1D), and we could identify two populations of IgG-SCs, one producing approximately 10-20 IgG/s (IgG^low^) and another one >200 IgG/s (IgG^high^). The frequency of IgG^high^-SCs was high in the spleen early in the immune response and decreased on day 14. In contrast, the frequency of bone marrow IgG^high^-SCs was more stable, and the separation between the subgroups more pronounced. Regarding affinity maturation, the slope distributions collapsed first in the spleen, i.e., demonstrated lower affinity on day 3, followed by an increase in affinity after that (Figure 1E). In the bone marrow, a significant shift towards higher affinities over time was observed (Figure 1E). Accordingly, our applied immunization protocol induced activation and differentiation of B cells into IgG-SCs.

**Figure 1:**
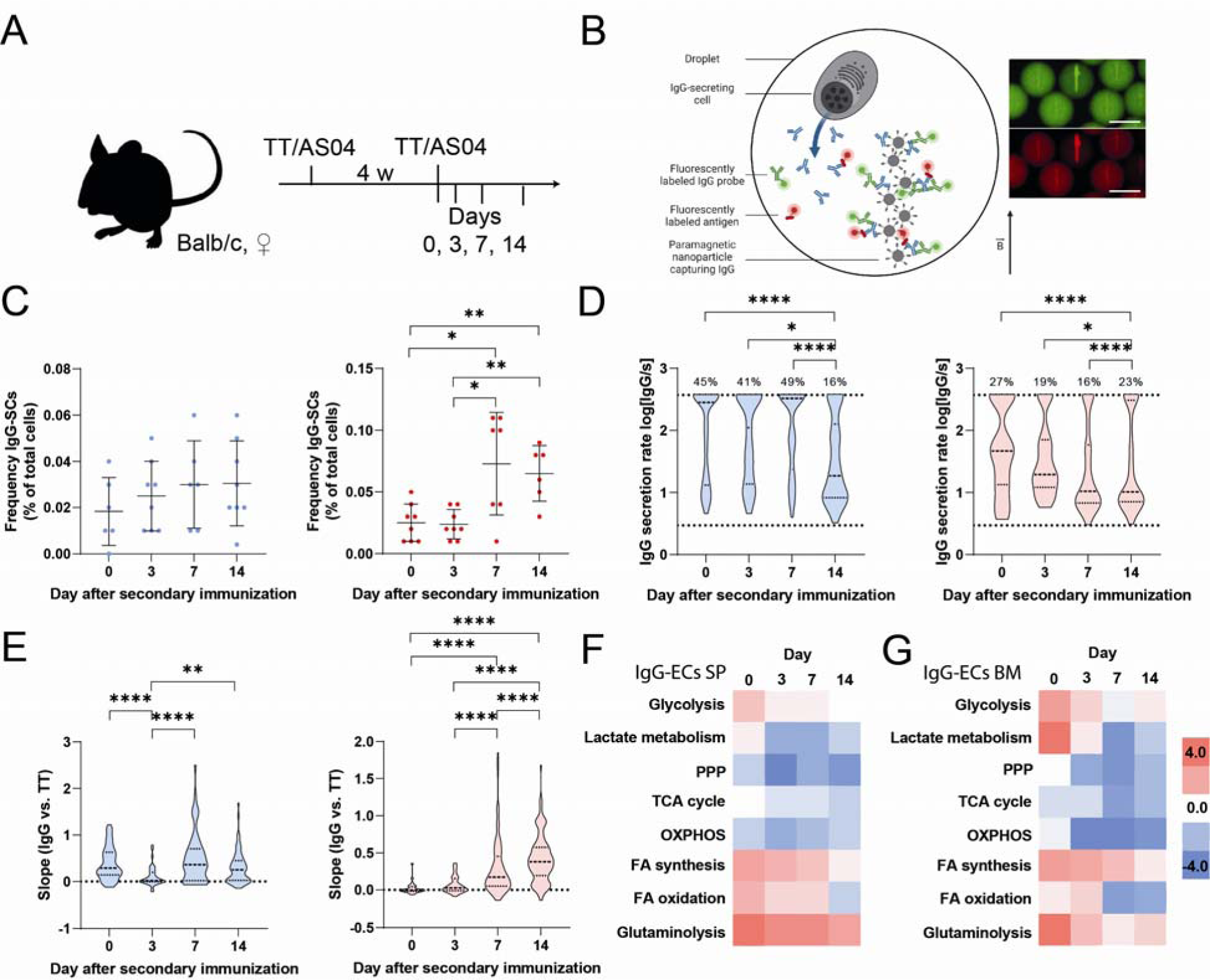
Summary of the induced immune response. (a) Employed immunization scheme and days of analysis. (b) Schematic representation of the bioassays to assess IgG secretion and affinity. The IgG secretion and affinity bioassay was a relocation sandwich immunoassay consisting of paramagnetic nanoparticles that form an elongated object in a magnetic field. Fluorescence images of the bioassays after 50 min are shown. Scale bar 50 μm. (c) Frequency of IgG-SCs (4 mice with measurements in the presence and the absence of the lactate assay, i.e., n_measurements_= 8, Grubb’s test was used to identify outliers). Cells with an IgG secretion rate equal or above the LoD (≥3 IgG/s) were defined as IgG-SCs. (d) The distribution of IgG secretion rates of IgG-SCs in the spleen (blue, left) and bone marrow (red, right) during the immune response. Data from four mice were pooled for the respective anatomical compartment and day (n_cells_= 187, 46, 96, and 82 in the spleen on days 0-14, n_cells_= 35, 37, 140, and 133 in the bone marrow on days 0-14). The percentage above the dotted line indicated the frequency of cells with an IgG secretion rate equal or above the assay’s quantitative range (≥375 IgG/s). (e) The distribution of affinity slopes of IgG-SCs in the spleen (blue, left) and bone marrow (red, right) during the immune response. Data from four mice were pooled for the respective anatomical compartment and day (n_cells_= 85, 40, 56, and 74 in the spleen on days 0-14, n_cells_= 28, 30, 140, and 120 in the bone marrow on days 0-14). (f and g) Analysis of gene expression changes of DEGs (average log 2-fold change) of selected metabolic pathways in IgG-ECs of the (e) spleen and (f) bone marrow on days 0, 3, 7, and 14. For the gene markers used to identify IgG-ECs, please refer to SITable 1. Cells from two mice were pooled in a ratio of 1:1 prior to scRNA-Seq, and sequences in the range of 800-5’500 cells were analyzed. Legend: pentose phosphate pathway (PPP), tricarboxylic acid (TCA) cycle, oxidative phosphorylation (OXPHOS), fatty acid (FA) synthesis, and FA β-oxidation (FA oxidation). For definition of glycolysis, lactate metabolism and TCA cycle see SIFigure 3. The level of statistical significance is denoted as *p <0.05, **p <0.01, ***p <0.001 and ****p <0.0001. Panel B was created with BioRender.com.

In parallel to the functional characterization of the IgG-SCs, a selection of metabolic pathways was investigated throughout the immune response by analyzing a subpopulation of the sampled cells with single-cell RNA sequencing (scRNA-Seq) and differentially expressed gene (DEG) analysis (p-value <0.05). First, we identified the population of plasma cells (PCs, SITables 1 and 2, and SIFigure 2) [50], and within the PC population, we identified the IgG-expressing cells (IgG-ECs), which correspond closest to the phenotypic IgG-SCs (see also methods *’scRNA-Seq - Pre-processing and processing of scRNA-Seq data’* and SITable 1). Interestingly, the frequency of phenotypic IgG-SCs and transcriptome-identified IgG-ECs did not correspond (see SITable 3), as around 10 times higher frequencies were found in the transcriptomic data set.

Compared to all other B cells of the samples, the DEG analysis revealed a significant upregulation of genes involved in glycolysis in IgG-ECs, with the strongest upregulation on day 0 in both spleen (Figure 1F) and bone marrow (Figure 1G). Most metabolic pathways displayed similar evolution in the spleen and bone marrow IgG-ECs. Genes involved in glutaminolysis, FA synthesis, FAO and glycolysis remained highly expressed over the early immune response to the booster. Most intriguing, we also observed a profound decrease in lactate metabolism, i.e., downregulation of the lactate dehydrogenase encoding gene (*LDHA*) in the immune response induced by the booster compared to day 0.

### Simultaneous measurements of lactate and antibody secretion rates do not alter the extracted functional data

While previous studies reported metabolic requirements of IgG-SCs [38], we wondered about the correlation between lactate metabolism, cellular functionality and survival, specifically since the GCs have been described as hypoxic in mature stages [22, 27]. Therefore, our next step was the introduction of a method to quantify LSRs of individual B cells into our droplet-based platform.

In literature, droplet-based systems have already been described to quantify lactate secretion of metabolically highly active cancer cells [51, 52]. Here, we adapted the assay to measure the LSRs of IgG-SCs and other splenic B cells accurately and precisely (Figure 2A, B). Therefore, we adapted a commercial lactate assay (see methods, SIFigure 4). In this assay, lactate secretion results in the enzymatically catalyzed substrate conversion into a fluorescent product, whereby the fluorescence intensity of the droplet was quantitatively correlated with the lactate concentration. First, we calibrated the sensitivity of our adapted assay (SIFigure 1). As we aimed to measure lactate secretion with a 10 min interval in-between measurements (as previously done for antibody secretion [48]), the calibration and additional control experiments revealed a quantifiable range of 0.10-0.80 amol/s for cellular LSRs (see also SI). Prior to the secondary immunization, the extracted LSRs for splenic cells from the B cell lineage were in the range of our calibration (0.18 ± 0.31 amol/s, Figure 2B), and 92% of all encapsulated B cells showed LSRs within the quantifiable range (Figure 2C).

**Figure 2:**
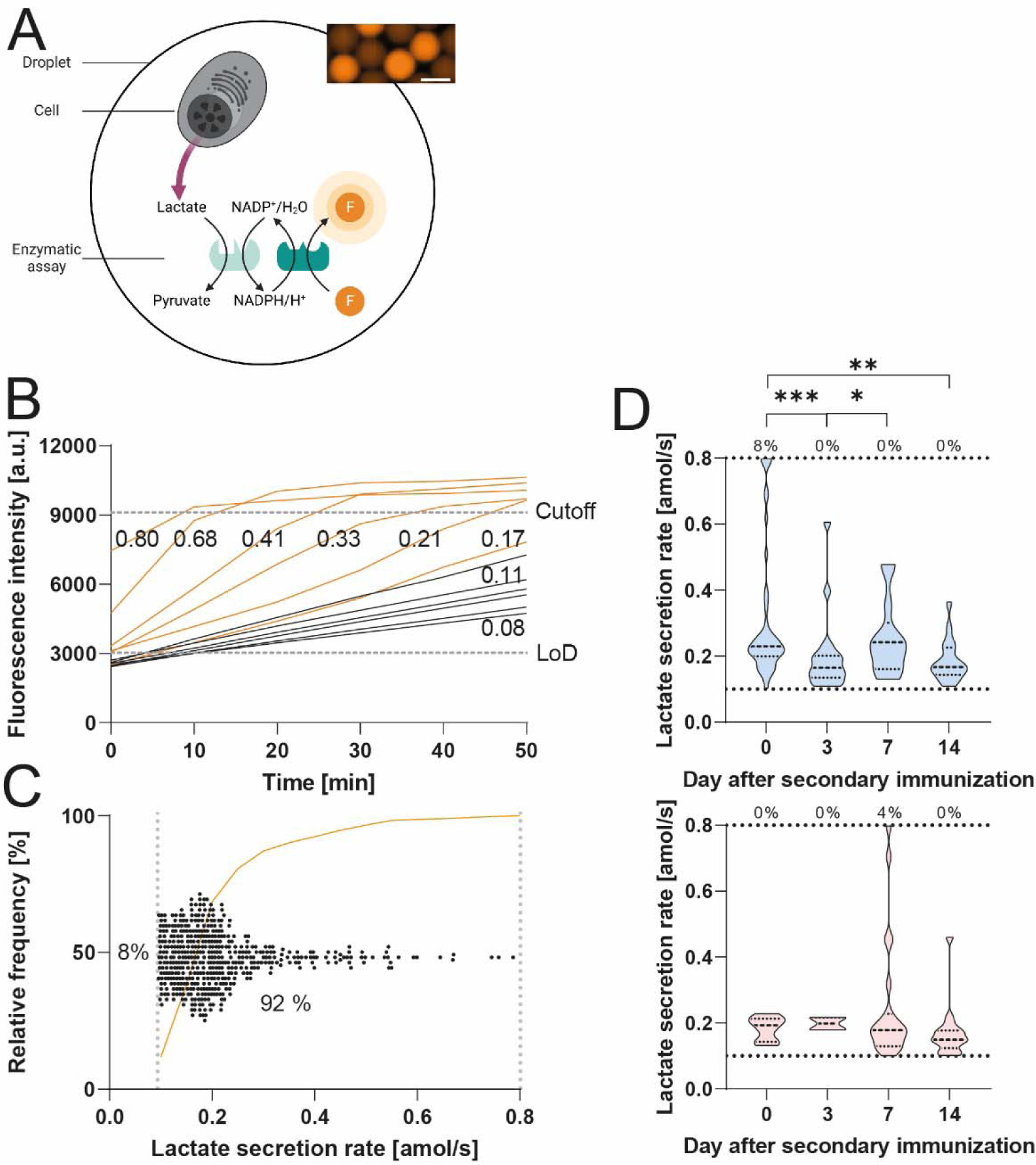
Lactate bioassay and cellular measurements. (a) Schematic representation of the bioassay to assess lactate secretion. The lactate assay consisted of an enzymatic kit resulting in a fluorescent product in proportion to the secreted lactate. A fluorescence image of the bioassay after 50 min is shown. Scale bar 50 μm. (b) Traces from droplets containing an individual B cell (orange, n= 6) and droplets without a cell (black, n= 6) and the corresponding LSRs in [amol/s]. The LoD and cutoff are shown. (c) The secretion rate of 500 randomly selected B cells was obtained from four mice on day 0, whereby 92% of cells displayed a secretion rate within the quantitative range. Every dot represents a cell, the cumulative frequency is shown, and the dotted grey lines represent the LoD and cutoff value. (d) LSRs of IgG-SCs in the spleen (blue, top) and bone marrow (red, bottom) during the immune response. Data from four mice were pooled for the respective anatomical compartment and day. The percentage above the dotted line indicated the frequency of cells with a LSR equal or above the assay’s quantitative range. n_cells_ in the distributions in (d) are between 13 and 51 in the spleen and 2 to 23 in the bone marrow. The level of statistical significance is denoted as *p <0.05, **p <0.01, ***p <0.001 and ****p <0.0001. Panel A was created with BioRender.com.

First, we investigated whether concomitant measurement of lactate secretion would alter the assessed cellular functions, the frequency of IgG-SCs, IgG secretion rate and affinity. For this purpose, these parameters were measured in the presence and absence of the lactate bioassay. In short, we did not find any significant difference in the assessed parameters in the combinatorial analysis (SIFigure 5). First, the obtained frequencies of IgG-SCs were normalized to the ELISpot results. The normalized frequencies obtained in presence and absence of the lactate bioassay differed non-significantly for the individual days and anatomical compartments, showing that simultaneous measurement and encapsulation did not alter the bioassay’s extracted functional data.

After demonstrating that the combination of lactate secretion, IgG secretion, and affinity measurements did not alter the respective data, the relationship between these parameters was investigated in IgG-SCs isolated from the spleen and bone marrow of immunized mice (n= 4 per time point). When looking at the LSRs of IgG-SCs over time, we observed that the LSRs of the IgG-SCs unified and decreased along the secondary immune response in the spleen (0.23 ± 0.20 amol/s on day 0 to 0.17 ± 0.06 amol/s on day 14, Figure 2D, p-value 0.003). In contrast, the median LSRs in the bone marrow IgG-SCs were rather stable (median LSR between 0.15-0.19 amol/s), with cells displaying the highest LSR range on day 7 (Figure 2D). Interestingly, the increase in the range of LSRs in the bone marrow correlated with the potential accumulation of affine IgG-SCs in this compartment (Figure 1C and E).

### Lactate secretion rates of IgG-SCs during an immune response do not correlate with the IgG secretion rate and affinity

To evaluate a potential relationship between lactate and IgG secretion rates at the single-cell level, IgG-SCs were first divided into high-secreting (>200 IgG/s) and low-secreting cells (3-200 IgG/s). No correlation between lactate and IgG secretion rates was found on individual days for the spleen and bone marrow (SIFigure 6). In addition, no correlation was observed at the single-cell level. Therefore, we concluded that cellular secretion rates for IgG and lactate were independent.

This relationship was further investigated by grouping IgG-ECs into IgG^low^- and IgG^high^-ECs (log2 fold change ≤1 for low *IGHG* expression) and comparing expression differences of *LDHA* (lactate metabolism) and other genes encoding for enzymes of key metabolic pathways (Figure 3). Gene regulation of the key metabolic pathways was generally similar between splenic IgG^low^- and IgG^high^-ECs (Figure 3A). Lactate metabolism was similarly activated in IgG^low^-and IgG^high^-ECs of the spleen on day 0, and both cell populations down-regulated the pathway subsequently with similar dynamics. In contrast, low IgG^low^- and IgG^high^-ECs differed in the expression level of glycolysis- and tricarboxylic acid (TCA) cycle-associated genes on day 3, and the pathways were only activated in the IgG^high^-ECs. In the bone marrow, the upregulation of *LDHA* expression was restricted to the IgG^high^-ECs on day 0 (Figure 3B). Subsequently, the IgG^high^-ECs showed decreased lactate metabolism activity compared to IgG^low^-ECs.

**Figure 3:**
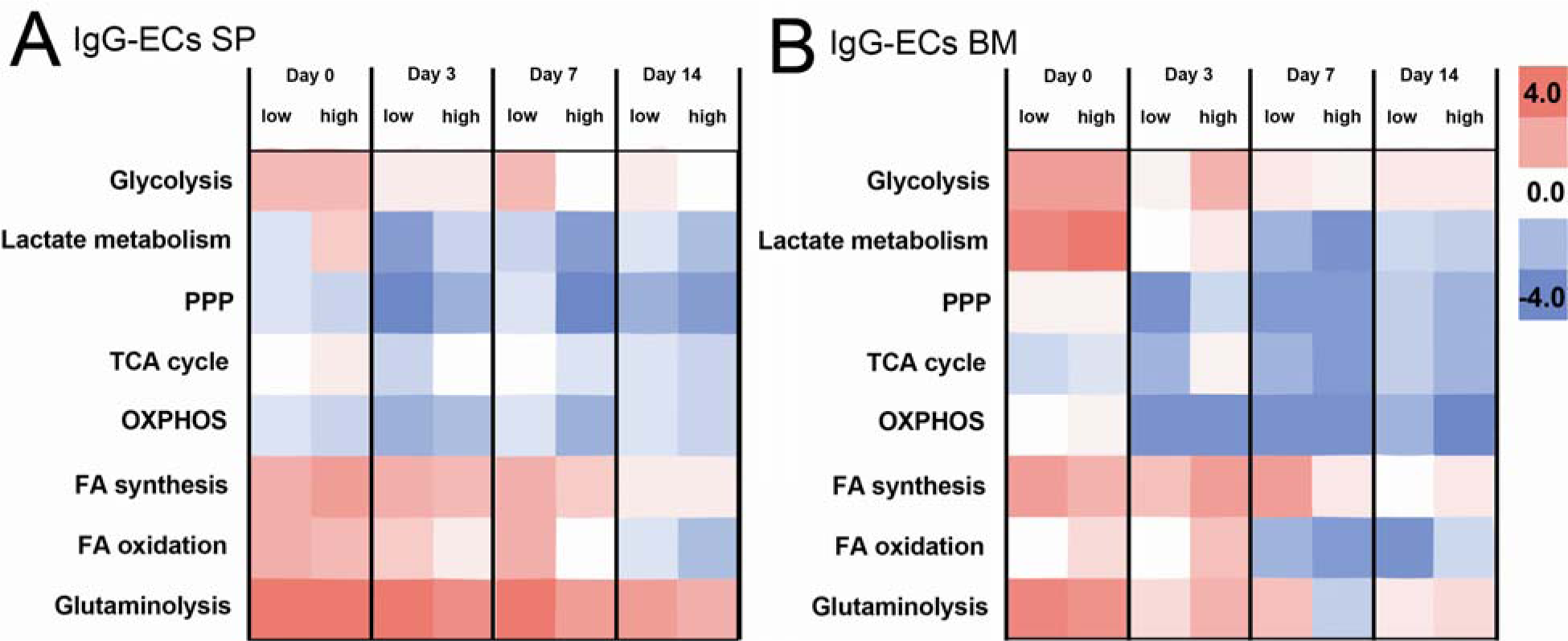
Comparison of the most relevant metabolic pathways and how their activity changed during the immune response based on gene expression changes of DEGs (average log 2-fold change) in IgG-ECs exhibiting low and high expression levels of IGHG in the (a) spleen and (b) bone marrow. IgG-ECs were categorized based on their IGHG expression level (threshold log2 FC 1), whereby 800-5’500 cells were analyzed.

Next, we explored the potential correlation between the affinity of the secreted IgG and LSR. IgG-SCs were divided into antigen-specific and non-specific cells (see method section *’Droplet-based measurements - assessment of antigen binding strength’*). We observed no significant differences between the two populations regarding their distribution of LSRs (SIFigure 6, p-value 0.92 and 0.60 for spleen and bone marrow, respectively), and no correlation at the single-cell level. Accordingly, cellular LSRs did not correlate with affinity.

### Lactate secretion is linked to the IgM isotype

We next performed additional measurements with an IgM probe on day 3 to investigate the potential relationship between lactate secretion and antibody isotypes. Interestingly, the splenic IgM-SCs (defined as cells with secretion rates ≥7 IgM/s, SIFigure 4) showed significantly increased LSRs compared to the IgG-SCs (0.29 ± 0.13 amol/s for IgM- and 0.20 ± 0.12 amol/s for IgG-SCs, respectively, p-value 0.0002, Figure 4A). The mean LSR of bone marrow IgM-SCs (0.26 ± 0.09 amol/s) was in the range observed for bone marrow IgG-SCs over the studied period of the immune response. Strikingly, all transcriptionally investigated metabolic pathways of splenic IgM-ECs (defined as *IGHM*-expressing PCs, SITable 1) were activated on day 3 compared to all splenic B cells, based on the average fold-change of gene expression levels (Figure 4B). In particular, lactate metabolism was prominently, but shortly, activated in IgM-ECs. In the direct response to antigen re-exposure (days 3 and 7), glycolysis, glutaminolysis, and FA synthesis-related gene expressions were mainly upregulated. On day 14, the cells focused on glutaminolysis while downregulating glycolysis- and FA synthesis-associated genes. In contrast to the highly dynamic behavior of splenic IgM-ECs, bone marrow IgM-ECs showed a relatively balanced activation of the pathways on days 0 to 7, except for glutaminolysis, which was activated (Figure 4C).

**Figure 4:**
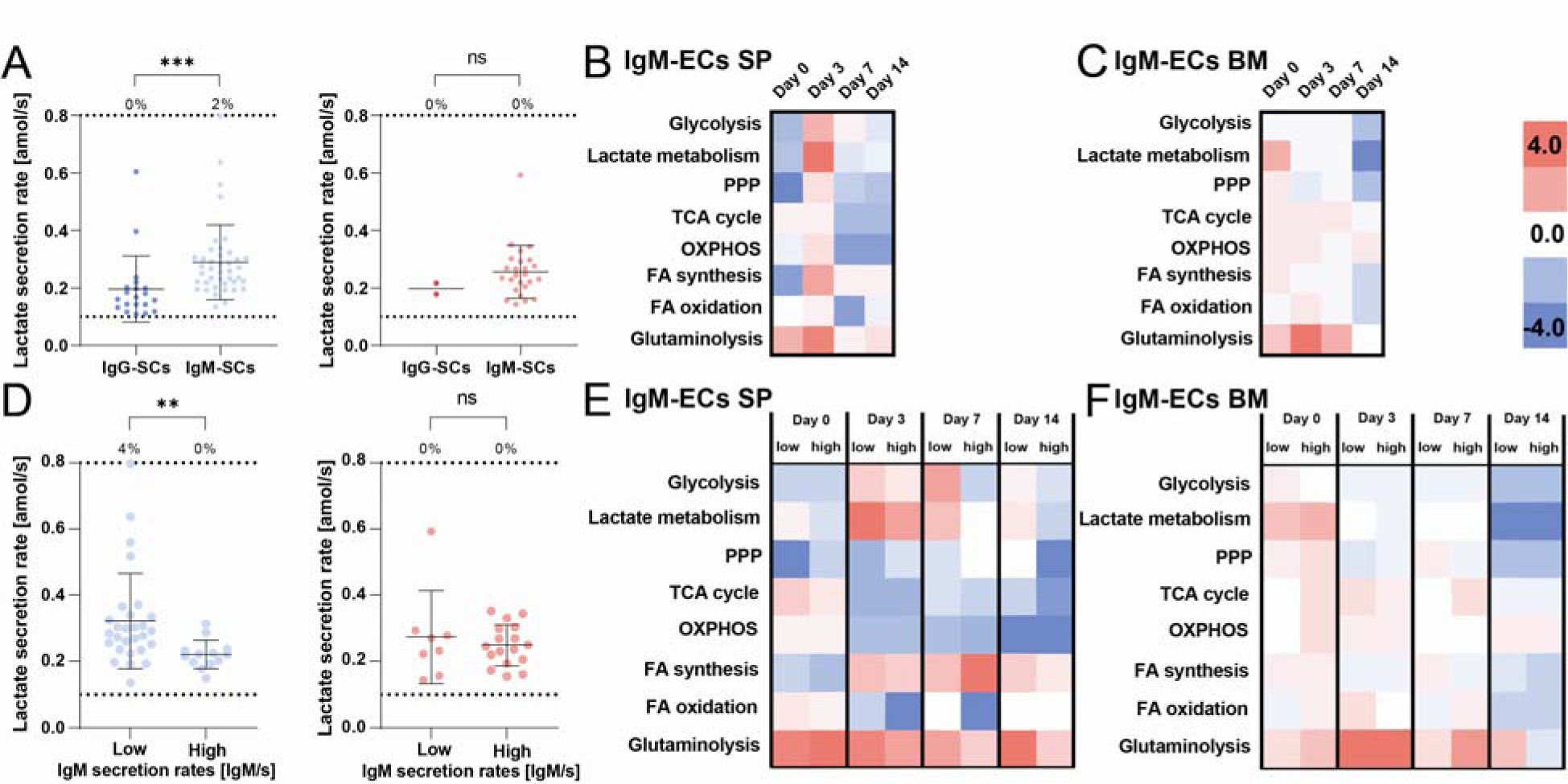
(a) Comparison of LSRs of IgG- and IgM-SCs of the spleen (blue, left, n_cells_= 20 and 41, respectively) and bone marrow (red, right n_cells_= 2 and 25, respectively). LSRs of individual cells, represented by a dot, are displayed. The percentage above the dotted line indicated the frequency of cells with a LSR equal or above the assay’s quantitative range (n_cells_= 20 IgG-SCs, 41 IgM-SCs for spleen, n_cells_= 2 IgG-SCs, 25 IgM-SCs for bone marrow). (b and c) Comparison of metabolic pathways activities of IgM-ECs of the (b) spleen and (c) bone marrow during the immune response. (d) Distribution of the LSRs between IgM^low^- and IgM^high^-SCs in the spleen (blue, left) and bone marrow (red, right). IgM-SCs were divided based on their IgM secretion rate (threshold 100 IgM/s, n_cells_= 28 high, 13 low for spleen, n_cells_= 8 high, 17 low for bone marrow). (e and f) Comparison of metabolic pathways activities between IgM^low^- and IgM^high^-ECs of the (e) spleen and (f) bone marrow. IgM-ECs were divided based on their IGHM expression level (threshold log2 FC 1). The level of statistical significance is denoted as *p <0.05, **p <0.01, ***p <0.001 and ****p <0.0001. Scale bars for B, C, E, and F are displayed in the figure.

Next, similar to the IgG-SCs, the IgM-SCs were divided into low- and high-secreting cells (threshold 100 IgM/s), and their LSR distributions were compared. Interestingly, we observed that the LSR distribution of IgM^low^-SCs was significantly shifted to higher rates in the spleen (Figure 4D, p-value 0.004) but not in the bone marrow (p-value 0.98). However, while the pooled data showed increased LSRs for IgM-SCs, no correlation was observed in either compartment at the single-cell level. We also compared the transcript levels of selected metabolic pathways between IgM^low^- and IgM^high^-ECs (see method and SI). Indeed, transcriptomic analysis confirmed an upregulated gene expression for lactate metabolism in both populations, but specifically in splenic IgM^low^-ECs on day 3 (Figure 4E). In the bone marrow, the IgM^high^-ECs generally displayed a higher gene upregulation of the studied metabolic pathways on day 0, including lactate metabolism, and later similar gene expressions as IgM^low^-ECs (Figure 4F). Therefore, the absence of correlation between LSRs and Ig secretion rates might only be linked to the IgG-SCs, although the data for IgM-SCs and -ECs was ambivalent.

### T cell-dependent response increases lactate secretion rates, whereby the addition of MPLA to the adjuvant system affects the magnitude

As we hypothesized that the observed dynamics in LSRs in IgG- and IgM-SCs were due to a direct response to the immunization component(s) or to immunization-induced environmental changes such as hypoxia and present cytokine repertoire, we further investigated the distribution of LSRs of all splenic B cells throughout the immune response. After antigen re-encounter, a substantial subpopulation with high LSRs (≥0.8 amol/s) was observed (Figure 5A), with a frequency of 27% at its peak. This subpopulation increased from day 3 to 7, collapsing on day 14. Contamination by other cell types was excluded, as flow cytometry confirmed a B cell lineage purity of 95 ± 2% (n= 7, see SIFigure 7).

**Figure 5:**
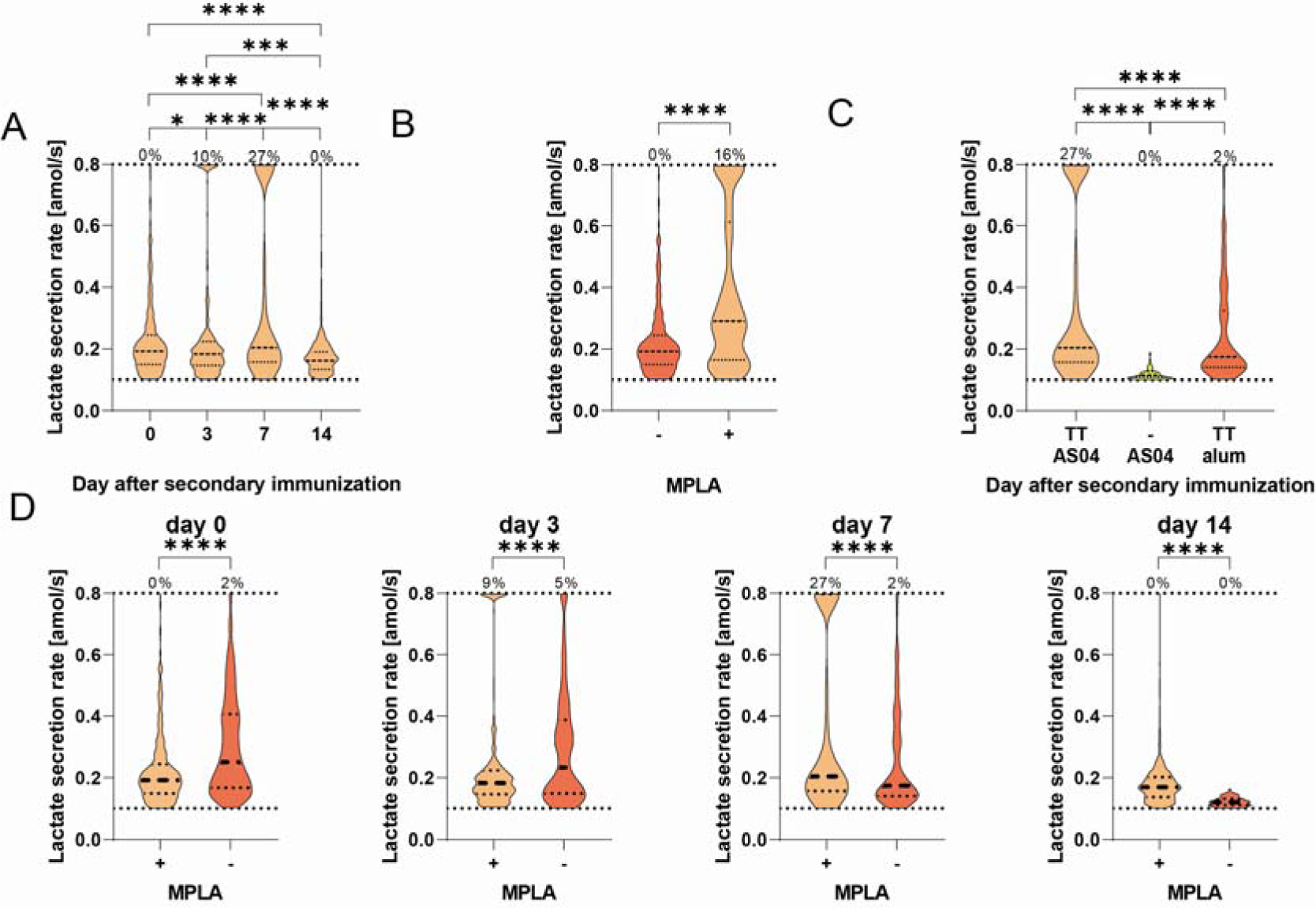
(a) Distribution of LSRs of all splenic B cells encapsulated during the measurements after immunization with TT/AS04. The secretion rate of 500 randomly selected B cells was obtained from four mice per day. The frequency above the dotted line indicated the frequency of cells with a LSR equal or above the assay’s quantitative range. (b) Ex vivo stimulation of splenic B cells harvested on day 0 with MPLA (1 μg per 1 x 10^6^ cells/mL, 37 °C, 4 h) resulted in a significant shift to higher LSRs (-: untreated cells, +: stimulated cells). The LSRs of 500 randomly selected B cells obtained from four mice per condition is displayed. (c) Comparison of LSRs of splenic B cells on day 7 after two immunizations with TT/AS04, AS04 (i.e., without antigen), and TT/alum (i.e., without MPLA). A total of 500 B cells were randomly selected per condition for comparison. (d) The distributions of LSRs from splenic B cells induced by immunization with TT/AS04 (indicated as + MPLA) or TT/alum (- MPLA) were significantly different over the observed timespan of the immune response. While TT/alum resulted in a distribution of secretion rates spanning the quantitative spectrum, TT/AS04 resulted in two populations, one with moderate and one with secretion rates ≥0.8 amol/s. The level of statistical significance is denoted as *p <0.05, **p <0.01, ***p <0.001 and ****p <0.0001.

As TLR4 signaling has been demonstrated to induce a shift towards glycolysis [7], we examined this possibility by *ex vivo* stimulation of splenic B cells harvested on day 0 with MPLA. As expected, the *ex vivo* stimulation led to a significant 1.5-fold increase in the LSRs (Figure 5B, p-value <0.0001), from 0.19 ± 0.11 amol/s to 0.29 ± 0.25 amol/s. Additionally, a population with high LSRs comparable to those observed on days 3 and 7 appeared after stimulation (from 0% to 16% after stimulation).

We were interested in the contribution of TLR4 and BCR signaling; therefore, we immunized mice with TT/AS04 (BCR and TLR4), TT/alum (BCR), and AS04 (TLR4). The T cell-independent immune response to AS04 alone led to a low median LSR (0.11 ± 0.02 amol/s) and a homogeneous population, suggesting that a T cell-dependent response is necessary for elevated LSRs and a heterogeneous distribution thereof on day 7 (Figure 5C). Moreover, the distribution of LSRs in mice immunized with TT/alum displayed a significant difference to distribution in TT/AS04 immunized mice (Figure 5C, p-value <0.0001). Thereby, cells displayed LSRs distributed over the quantitative range, and no distinct subpopulation with high secretion rates (≥0.8 amol/s) was observed. As the dynamics of LSRs might be shifted due to temporal differences in the immune responses, LSRs of splenic B cells of mice immunized with TT/alum were further analyzed over time (Figure 5D). While all distributions were significantly different compared to the TT/AS04 immunizations (p-value <0.0001 for all days), the observations were not biased by different immune dynamics, i.e., immunizations with TT/alum resulted in very few LSR^high^ cells. Therefore, neither the antigen nor MPLA alone was responsible for the appearance of these LSR^high^ cells from the B cell lineage.

### High lactate secretion rates relate to altered pH homeostasis and apoptosis in splenic B cells

As the immunization with TT/alum led to considerably higher frequencies of IgG-SCs (SIFigure 8), we wondered whether this was related to the lower accumulation of cells with LSRs of ≥0.8 amol/s. Therefore, we investigated the relationship between LSRs and cellular homeostasis, namely oxidative stress, pH homeostasis, and apoptosis.

A cell-permeable, ROS-sensitive reagent was added to the cells to assess the ROS production level of splenic cells (Figure 6A, SIFigure 9 for controls with ROS scavenger and single-cell data). Different subpopulations were observed on day 0, and the subpopulations with a higher ROS level diminished significantly throughout the response to secondary immunization (distribution differed significantly between all days, all p-values <0.0001). We categorized the B cells into ROS^high^ and ROS^low^ subgroups (see methods, SIFigure 9) and compared their distribution of LSRs throughout the immune response (Figure 6B). On day 0, ROS^high^ cells displayed higher LSRs (p-value 0.0008), with LSR^high^ cells in both populations. Differences were still visible on day 3 (p-value 0.03) but disappeared afterward.

**Figure 6:**
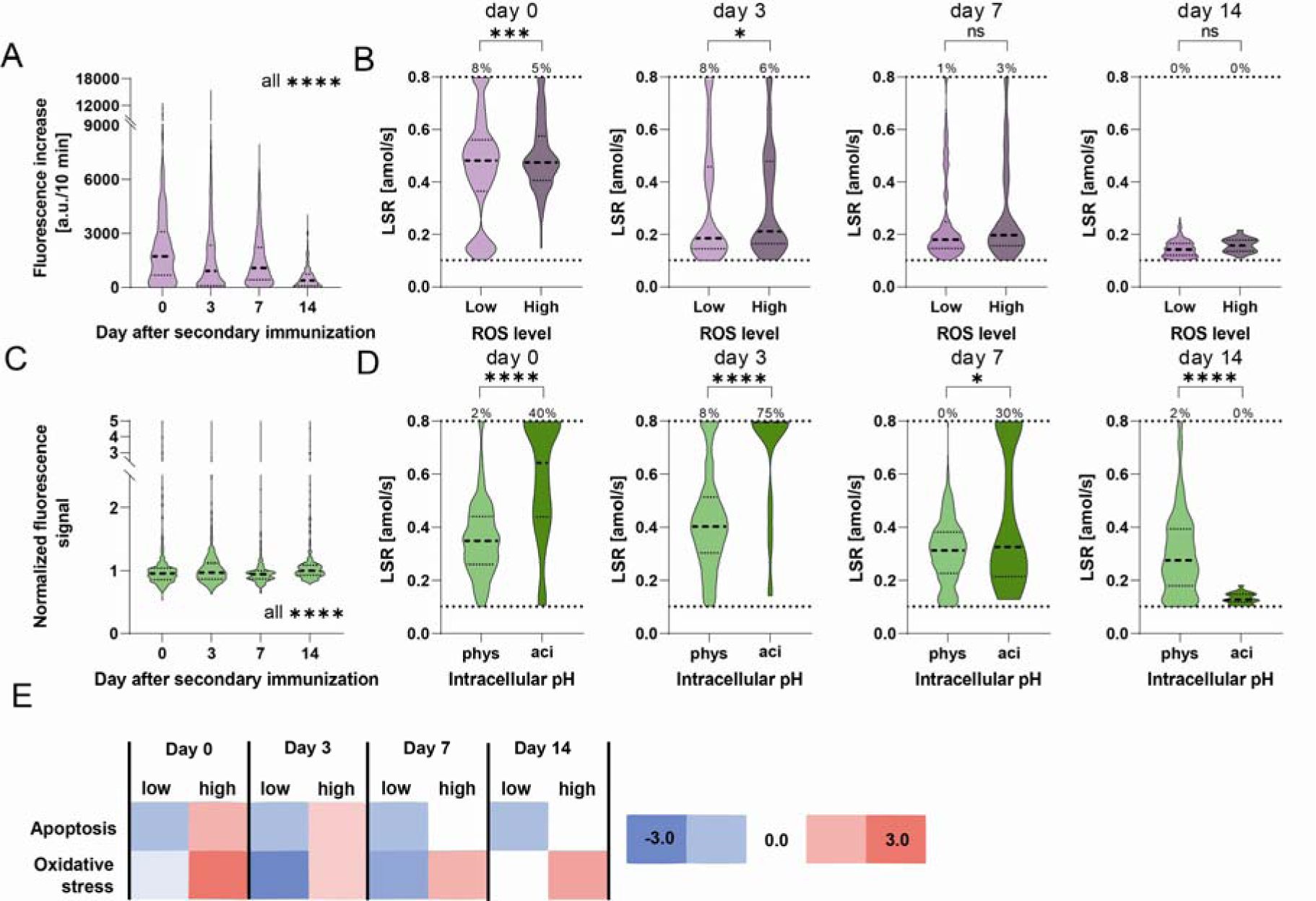
(a) Distribution of the increases in cellular fluorescence intensities, which served as an indicator of the cellular ROS production level of splenic B cells during the immune response (in purple). A random selection of 125 cells per mouse (4 mice) was pooled, and the distributions were significantly different from each other on all days (p <0.0001). (b) Comparison of LSR distributions of the same cells categorized into ROS^low^ and ROS^high^. The frequency above the dotted line indicated the frequency of cells with a LSR equal or above the assay’s quantitative range. (c) Distribution of the normalized fluorescence signal of splenic B cells indicating the pH_i_ throughout the immune response (in green). A random selection of 125 cells per mouse (3 mice) was pooled, and the distributions were significantly different from each other on all days (p <0.0001). (d) The cells were categorized according to their pH_i_ into cells with physiological (phys) or acidic (aci) pH_i_, and their distributions of LSRs were compared. The number above the dotted line indicated the frequency of cells with a LSR equal or above the quantitative range on the respective day. (e) Correlation between lactate metabolism and expression level of genes associated with apoptosis and oxidative stress in splenic B cells throughout the immune response. Cells were categorized according to lactate dehydrogenase (LDH) encoding gene expression into ‘low’ and ‘high’. The level of statistical significance is denoted as *p <0.05, **p <0.01, ***p <0.001 and ****p <0.0001.

To assess the relationship between LSRs and pH homeostasis, the cells were stained with a pH-sensitive dye, and the normalized cell’s fluorescence intensity was used as intracellular pH (pH_i_) indicator (see methods, SIFigure 10). Figure 6C shows the distributions of the normalized cell signals of the splenic B cells studied during the immune response (significant difference between all days, all p-values <0.0001), and Figure 6D upon dividing cells into two groups with physiological and acidic pH_i_ (see also methods). Interestingly, we found a correlation between acidic pH_i_ and high LSRs on days 0-7, whereby cells with acidic pH_i_ displayed significantly higher LSRs (p-values <0.0001, <0.0001 and 0.03, respectively). This finding indicates that lactate generation exceeded the cellular transport capacities for lactate and protons on these days.

As the acidic pH_i_ cell state correlated with LSR^high^, we next aimed to correlate LSRs to apoptosis. First, we employed a bioassay to investigate whether caspase-3 and -7 were present in their active forms, which are involved in a potential apoptosis pathway of B cells. Surprisingly, caspase-3/7 activity was correlated with lower LSRs (see SIFigure 11). However, caspase-3 has also been described to play a vital function in B cell proliferation [53] and, thus, might not indicate apoptosis in this case. However, when analyzing the transcriptomic data, we found that expression of genes associated with oxidative stress and apoptosis were upregulated in high lactate-metabolic B cells on days 0 and 3 (Figure 6E). Afterward, apoptotic genes were still upregulated in high lactate-metabolic B cells but to a reduced level, and the expression level of oxidative stress genes remained similarly upregulated on both days.

## Discussion and conclusion

Various links between metabolic programs and plasma cell differentiation and function have recently been reported [25, 29-31, 46, 54, 55]. However, studies were limited to indirect assessments of metabolism by studying expression levels of genes or proteins, static measurements of metabolites or substrate uptake, or the direct measurement of metabolic fluxes at the population level. In contrast, a droplet-based technology allowed the analysis of LSR of cancer cells at the single-cell level. Thereby, cells were individually encapsulated into droplets, which served as independent reaction vessels, and by immobilizing, they observed secretion dynamically [48, 49]. In this study, the parallel dynamic measurement of LSR and functional parameters of B cells was developed. We used single-cell transcriptomic analysis to add cellular context and information about alternative key metabolic pathways. The integrated functional and metabolic assays provided a starting point to characterize key metabolic changes present in B cells at various stages on the single-cell level. Indeed, we identified a crucial need for further development of technologies for the direct and parallel measurement of metabolic fluxes and cellular functionalities linked to identity to better understand their complex relationships [38]. Here, measurements of the LSR was combined with functionality measurements, namely IgG and IgM secretion rates and affinities of the secreted antibodies, as well as with measures of cellular homeostasis indicators, namely ROS production level, intracellular pH, and caspase-3/7 activity.

For the study presented here, we focused on B cell activation and differentiation into IgG-SCs after re-immunization with TT/AS04. However, by concentrating on re-activating previously formed B cells, we also introduced certain relics of the previous immunization into our study. Considering the collapse of the affinity slope distribution on day 3 in the spleen, a cellular turnover between days 0 and 3 was concluded. Accordingly, the cells resident in the spleen late after primary immunization emigrated, stopped secreting, or died. Migration to the bone marrow was unlikely due to the absence of a concurrent increase of IgG-SCs. Furthermore, the affinity reduction of splenic IgG-SCs on day 3 was surprising as an extrafollicular B cell response was expected, whereby B cells with higher affinity and IgG memory cells resulting from the primary response were predisposed to this cell fate [56, 57]. One reason could be that the primary response was still in its effector phase on the day the booster was performed, although GC dissolution was reported after day 21 [58, 59]. In contrast, Pettini and colleagues have shown that subcutaneous immunizations supplemented with alum lead to immune responses in the effector phase after four weeks in spleen and iliac lymph nodes [60]. Their observation was in line with our histology observations that GCs were present early after secondary i.p. immunization (unpublished data). In summary, our study did not only investigate a pure recall immune response upon antigen re-encounter but also the influence of the still ongoing primary immune response.

Regarding our observation that GCs still appeared active on day 0 of our study, the IgG-SCs present in the spleen on day 0 could originate mainly from the GC response to the primary immunization, whereas the IgG-SCs on days 3 and 7 could rather result from extrafollicular reaction to antigen re-encounter. Hence, the observed lactate metabolism activation originated primarily in recalled IgG-ECs from the previous GC response. This connection was especially conceivable because GCs have been described as poorly vascularized, hypoxic environments [22, 27] and because hypoxia leads to the adaptation of cellular metabolism by increasing the conversion of pyruvate to lactate [22, 23, 61] and inhibiting pyruvate’s entry into the TCA cycle [23]. Secondly, a fast adaptation due to re-oxygenation of hypoxic cells might alter the metabolic fluxes rapidly when the cells were removed from the environment *in vivo*. However, we still were able to characterize differences between the days, indicating that at least some metabolic flux differences remained.

When characterizing the immune response after re-exposure in the spleen and bone marrow, we observed the transcriptional upregulation of key metabolic pathways in IgG-ECs, diminishing after booster immunization. By direct measurement, we found that the LSR of splenic IgG-SCs was dynamic, with a reduction of the median secretion rate along the immune response with the disappearance of LSR^high^ IgG-SCs. Interestingly, the direct flux measurements using droplets and the transcriptomic analysis did not evolve similarly, hinting toward potential post-transcriptional, translational, and allosteric regulations [42-44]. The absence of a correlation between lactate and IgG secretion led to the hypothesis that these cells could originate from hypoxic environments like GCs (day 0) or, additionally, that they have only recently been activated, whereby effects from activation-induced ROS may still be present (days 3 and 7). Both hypoxia and ROS were shown to lead to the stabilization of hypoxia-inducible factor (HIF)-1α [22, 62]. As HIF-1α was reported to reduce the level of high-affinity antibodies due to its inhibitory effect on mTORC1, which is essential for the expression of AID [63], we hypothesized that LSRs and affinity are negatively correlated. However, we did not find such a correlation in the data.

In contrast to the spleen, an increase in the frequency of IgG-SCs was observed in the bone marrow on days 7 and 14. This observation fits with findings of IgG-SCs in the blood one week after primary immunization [64, 65], which home to the bone marrow to compete for survival niches [66], and with reports of the appearance of ASCs newly formed after secondary immunization in the bone marrow from day 4 onwards [67]. Additionally, the frequency of IgG-SCs in purified B cells might be even higher, as the employed kit only partially depleted non-B cell lineage cells from the bone marrow samples. Furthermore, the observed shift in the distribution of the affinity slopes indicated the immigration of antigen-specific IgG-SCs. Based on the IgG secretion rate distribution, these newly immigrated cells appeared to be low IgG secretors with constant LSRs [68].

The LSR distributions changed throughout the immune response induced by TT/AS04, with a subpopulation secreting high amounts of lactate (≥0.8 amol/s) present on days 3 and 7. By immunizing mice with AS04 alone (i.e., without TT), we could show that a T cell-dependent immune response was required for increased LSRs on day 7. TLR4 signaling was shown to induce glycolysis [7], a finding also made in our *ex vivo* stimulation experiments. However, TLR4 signaling alone did not generate the observed dynamics and magnitude in LSRs during the immune response. While immunization with TT/AS04 and TT/alum increased median LSRs, their distributions of LSRs varied significantly over the immune response, and both had a subpopulation with LSRs^low^ in common. However, while the other cells of the TT/alum condition were distributed over the quantitative range, TT/AS04 induced a second distinct subpopulation with secretion rates ≥0.8 amol/s. As this subpopulation was absent on day 0, we conclude that the booster induced this phenotype. An explanation for their high secretion rate could be the combination of BCR and TLR4 signaling. Comparing the distributions of days 0 and 14, lactate generation appeared upregulated again during the sustained GC reaction, potentially due to hypoxia [22, 27].

The fact that high LSRs were found in the TT/AS04-condition which resulted in lower frequencies of IgG-SCs, made us wonder whether LSRs^high^ negatively affected cellular homeostasis and potentially led to cell death. Studies have shown that ROS played an important role in activating B cells [5, 69-75]. ROS present in B cells were either produced by NADPH oxidases [5, 74, 75] or as byproducts from the mitochondrial respiration chain and oxidative protein folding [5, 69-73]. Apart from the need of ROS for cell activation and signaling, ROS upregulation might mediate various consequences for the cell, such as oxidative stress and apoptosis [76]. In addition, metabolic enzymes, including the TCA cycle enzyme aconitase, were described as susceptible to inactivation by ROS [77], forcing cells to switch to alternative metabolic pathways.

Indeed, the LSR distributions of cells with high ROS levels were significantly elevated on days 0 and 3. Moreover, cells with high LSRs were present in both groups, indicating that high lactate generation was not necessarily linked to high cellular ROS levels. Moreover, the presence of cells with increased LSR was most pronounced on day 7, and on this day, the distribution of LSRs between cells with low and high ROS levels was non-significantly different. This led us to the conclusion that high LSRs and cellular ROS levels were not directly linked. Nevertheless, our transcriptomic analysis revealed the activation of oxidative stress in high lactate-generating cells [78]. The analysis further revealed that activation of lactate metabolism was accompanied by activation of glutaminolysis, a pathway also linked to the production of the antioxidant glutathione [79]. This observation was further consistent with findings that ROS increased antioxidant gene expression [80-82]. Collectively, our data showed a potential link between lactate metabolism and oxidative stress, but a direct correlation was not observed.

We identified a disturbance of cellular pH homeostasis as another effect of high lactate generation. Indeed, an accumulation of intracellular lactate might reduce pH_i_ [83], and B cells with an acidic pH_i_ showed increased LSRs on days 0 to 7. Accordingly, at least some B cells might not adapt to high glycolysis like cancer cells, which exploit proton pumps and transporters to obtain an alkaline pH_i_ despite high lactate generation [84]. Extracellular acidosis may further contribute to a reduction in pH_i_ [85]. One consequence of a possible intra- and extracellular accumulation of lactic acid could be the inhibition of proliferation and the promotion of cell death, as shown in T cells [86]. Congruently, *in silico* analysis of pH-dependent metabolism revealed that acidic pH_i_ impairs cell proliferation [87]. Our transcriptomic analysis revealed an activation of apoptosis in cells with increased lactate metabolism, supporting our observation of a correlation between LSR and acidic, potentially lethal pH_i_. These results supported the suggestion that LSR was related to the cellular environment and not simply dependent on antibody secretion rates and antigen specificity. In summary, we concluded that there was likely a link between LSR, intracellular pH, and apoptosis. Whether the observed link is a correlation or a causal relationship and the investigation of the underlying mechanism of this relationship require further studies.

## Acknowledgments

The European Research Council starting grant (grant #80’336), the Swiss National Science Foundation (grant #310030_197619), and the Branco Weiss - Society in Science Foundation supported this work.

## Contributions

O.T. M.B. planned all experiments, performed and analyzed the droplet experiments, and drafted the manuscript. D.R.performed and analyzed the scRNA-Seq experiments. K.P. was involved in adapting the functionality assay, and A.L. ran the ELISpot measurements. M.T. conducted and analyzed the flow cytometry measurements under the supervision of C.H. K.E. supervised the project. All authors commented on and revised the manuscript.

## Data availability

The transcriptomic data generated and analyzed during the current study will be publicly available on the ArrayExpress repository under E-MTAB-13205.

## Methods

### Mice immunization

BALB/cJRj (Janvier Labs, female, 7-10 weeks old at time of primary immunization) were immunized with recombinant tetanus toxin heavy chain fragment C (TT, FinaBio; 10 μg). All immunizations were performed intraperitoneally (i.p.) with 100 μL volume. For immunization, the antigen was added to physiological water (InvivoGen, vac-phy) and mixed with adjuvant system 04 (AS04: 5 μg MPLA-SM (InvivoGen, tlrl-mpla2) and alhydrogel adjuvant 2% (alum, InvivoGen, vac-alu-250)) or alhydrogel adjuvant alone in a ratio of 1:1 (*v/v*). The mice were re-immunized with the same mixture four weeks after primary immunization (termed day 0). The immune response was examined on the day of re-immunization (day 0, i.e., mice were not re-immunized) and 3, 7 and 14 days after re-immunization. Experiments were performed according to institutional guidelines and Swiss Federal regulations and were approved by the cantonal veterinary office Zurich (animal experimentation license ZH215/19).

### Cell isolation

On the day of measurement, the spleen and bone marrow of the mice were harvested. Spleens were dissociated by gently passing the tissue through a 40 μm cell strainer. To recover bone marrow cells, both tibias and femurs were flushed using sterile-filtered ice-cold MACS buffer (D-PBS #P4417 with added 2 mM EDTA #03620 and 0.5% BSA #A3059, all Sigma Aldrich). Afterward, spleens and bone marrow cell suspensions were collected by centrifugation (400 g for 5 min, 4 °C), and red blood cell lysis was subsequently performed at room temperature (1 min, BD Pharm lyse Lysing buffer). Cells were filtered (40 μm cell strainer) and washed twice with ice-cold MACS buffer. The cells were subsequently processed using the Pan B Cell Isolation Kit II (Miltenyi Biotec) according to the manufacturer’s protocol. The cells were stored on ice (≤12 h).

### Droplet-based measurements

*Droplet-based measurements - Aqueous phase I: Preparation of cells and lactate, IgG and IgM standards for droplet creation* Cells were stained with either CellTrace^TM^ Violet (Thermo Fisher, C34571), CellTrace^TM^ Far Red (Thermo Fisher, C34564), or pHrodo^TM^ Green AM (Thermo Fisher, P35373) at 2 x 10^6^ cells per mL. CellTrace^TM^ Violet (5 μM) and pHrodo^TM^ Green AM (10 μM, addition of 1X PowerLoad^TM^) stainings were performed in HBSS for 30 min on ice, CellTrace^TM^ Far Red (1 μM) staining in HBSS for 10 min on ice. After incubation, the cells were washed with ice-cold MACS buffer. The cells were collected (400 g, 5 min, 4 °C) and re-suspended in ice-cold assay buffer (RPMI 1640 without phenol red, #11835063, 10% (*v/v*) KnockOut Serum Replacement, #10828010, 1X penicillin-streptomycin, #10378016, 25 mM MOPS pH 7.5, #J61843, 0.1% (*v/v*) Pluronic F-127, #11835030, all Thermo Fisher, and 0.5% (*w/v*) recombinant human serum albumin, Sigma Aldrich, A9731) to achieve a λ (mean number of cells per droplet) of 0.25-0.50. Lactate standards (Lactate Assay Kit, Sigma-Aldrich, MAK064) with concentrations from 2-200’000 amol/nL were prepared. IgG (Kerafast, EFD006) and IgM isotype controls (Biolegend, 401601) of 100 nM were prepared, with eight serial dilutions by a factor 2.

### Droplet-based measurements - Aqueous phase II: Preparation of nanoparticles and detection reagents

A commercial fluorometric lactate assay (Lactate Assay Kit, Sigma-Aldrich, MAK064) was used to measure secreted lactate with slight adaptation in its composition. The enzyme concentration of the kit was increased to enhance the turnover rate due to higher lactate concentrations in droplets compared to well plates (4x of manufacturer’s concentration, i.e., 8 μL enzyme mix per 100 μL bead solution), whereas the concentration of the fluorescent probe was decreased due to photoactivation at higher concentrations (0.5x of manufacturer’s concentration, i.e., 0.5 μL fluorescent probe per 100 μL bead solution). Paramagnetic nanoparticles (Strep Plus, 300 nm, Ademtech) for measurement of IgG- and IgM-secretion were prepared as described elsewhere [48, 49]. The IgG assay contained a goat anti-mouse IgG Fc antibody (SouthernBiotech, 1033-31, labeled with Alexa Fluor 488) and the IgM assay a donkey anti-mouse IgM F(ab’)_2_ (Jackson ImmunoResearch, 715-476-020, labeled with DyLight 405), both at a final, in-droplet concentration of 75 nM. To assess the affinity, the antigen TT was labeled in-house with APC (APC Conjugation Kit – Lightning-Link®, Abcam, ab201807; according to manufacturer’s protocol) and added to obtain a final, in-droplet concentration of 50 nM. The caspase-3/7 activity assay contained CellEvent Caspase- 3/7 Green Detection Reagent (Thermo Fisher, C10723) at a final, in-droplet dilution of 1:1000 from the manufacturer’s stock. CellROX Green Reagent (Thermo Fisher, C10492) was added to indicate the ROS level in a final, in-droplet dilution of 1:800 from the manufacturer’s stock. The aqueous phase II solution was stored on ice (≤12 h) and re-suspended prior to measurements.

### Droplet-based measurements - Microfluidic droplet generator and observation chamber assembly

The production of microfluidic PDMS chips for droplet generation and the 2D observation chamber were produced as detailed in the references [48, 49].

### Droplet-based measurements - Cell stimulation

In addition to the direct droplet measurements, a sample of the splenic cells extracted on day 0 was *ex vivo* stimulated with MPLA (1 μg/1 x 10^6^ cells/mL) for 4 h at 37 °C. Afterward, the cells were collected and processed as described above.

### Droplet-based measurements - Data acquisition and analysis

Droplets of 50 pL were generated and transferred into the microfluidic observation chamber as previously described [48, 49]. The cell-containing aqueous solution I was cooled using ice-cold water during encapsulation. After complete filling of the observation chamber, the container was closed and mounted onto an inverted fluorescence microscope for imaging (Ti2 Eclipse, Nikon) equipped with a motorized stage, excitation light (Lumencor Spectra X) and a digital CMOS camera (ORCA-Fusion C14440, Hamamatsu, in 2x2 binning mode). Fluorescence signals for the specific channels were recorded using appropriate bandpass filters (DAPI, FITC, TRITC and Cy5 filter set, Semrock) at 37 °C and ambient oxygen and carbon dioxide concentrations. Images were acquired using a 10x objective (NA 0.45, Nikon). An array of 10x10 images, corresponding to around 50’000 droplets, were acquired for each experiment with a 10 min-interval over 50 min (6 measurements per experiment). The images were analyzed using an updated custom Matlab script (Mathworks, Versions 4.032 and 4.106, previous version published in references [48, 49]). Values were exported to Excel (Microsoft), and secretion rates and other metabolic and functional parameters were extracted as described below. Droplets were visually controlled, and droplets containing multiple cells were excluded from further analysis of metabolic parameters. The experiment-specific encapsulation parameter λ was determined using a previously published script [88]. Outliers, i.e., data from mice with extremely high responses, were excluded using Grubb’s test (p <0.05). For paired t-tests, the Wilcoxon matched-pairs signed rank test was used, and for the comparison of distributions, the Kolmogorov-Smirnov test (GraphPad Prism, Version 8.2.0). The level of statistical significance is denoted as *p <0.05, **p <0.01, ***p <0.001 and ****p <0.0001.

### Droplet-based measurements - Calibration curve for lactate

Droplets were generated and imaged as described above. The fluorescence signal of the whole droplet was determined. The calibration curve was fitted using a sigmoidal (4 PL) nonlinear fit, whereby x was the lactate concentration (GraphPad Prism Version 8.2.0). The detection limit (LoD) was determined based on an adaptation of the LoD definition by Armbruster and Pry [89]:

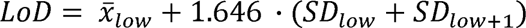

Where *x¯_low_* is the median of the 2 amol/nL calibration point, and SD is the standard deviation (SD) of the 2 amol lactate/nL (low) and 20 amol/nL (low+1). The cutoff, or upper limit of quantification, was calculated based on the 95% confidence interval (CI) of the sigmoidal fit of the calibration. For this purpose, the SD of the 95% CI was calculated, followed by the subtraction of 2x the SD of the plateau value (best-fit value):

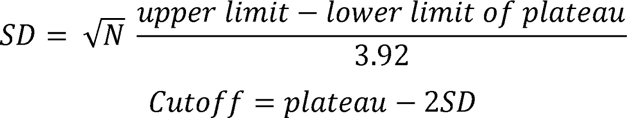

where N is the number of individual concentrations used for fitting and the upper and lower limit of the plateau result from the 95% CI. The quantitative range was adjusted due to some limitations and resulted in 0.10-0.80 amol/s (see SI section 2).

### Droplet-based measurements - Calculation of lactate secretion rate

The droplet intensities were converted to lactate concentrations using the calibration curve. The secretion rate of individually encapsulated cells was calculated by dividing the concentration change by the time interval, followed by the calculation of the average.

### Droplet-based measurements - Calibration curve for IgG/ IgM

Droplets were generated and imaged as described above. The ratio between the mean fluorescence signal on the nanoparticles and the mean fluorescence signal of the whole droplet was determined. The calibration curve was fitted using an exponential model according to Aymerich *et al*. [90]:

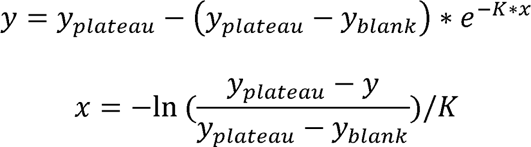

where x was the IgG/ IgM isotype control concentration, y was the fluorescence relocation, y_blank_ was the median relocation of a blank sample, and y_plateau_ was the fitted plateau value. To fit the data, a random selection of individual droplets for the different concentrations was used (around 50’000 droplets per concentration). To define the upper limit of the quantitative range of the assays, a cutoff was set on the fluorescence relocation equal to the plateau value from the exponential fit minus the half-width of the prediction interval (90% confidence level). The resulting quantitative ranges were 3-375 IgG/s and 7-150 IgM/s (see SIFigure 4). Cells with a secretion rate ≥3 IgG/s were defined as IgG-SCs and cells with a secretion rate ≥7 IgM/s as IgM-SCs.

### Droplet-based measurements - Assessment of IgG/ IgM secretion rates

The secretion rate of antibodies was calculated according to Aymerich *et al*. [90]. In short, the ratio between the mean fluorescence signal on the nanoparticles and the mean fluorescence signal of the whole droplet was determined and converted to an IgG/ IgM concentration based on the calibration curve. Droplets positive for the following three criteria were visually verified for secretion:

i. The droplet displayed at least one time point a relocation signal above the LoD.
ii. The relocation signal of the analyte increased over time.
iii. The signal varies more over the time series than the analytical noise in the absence of analyte:

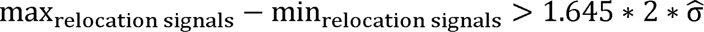

Whereby the definition of the LoD was:

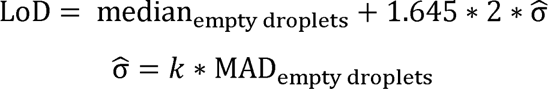

with MAD_empty droplets_ being the median absolute deviation in droplets without a cell (termed empty droplets), k = 1.4826, and k*MAD a consistent estimator of the standard deviation σ, considering the distribution of relocation signals for empty droplets as normal. The secretion rate (molecules/s) was determined by calculating the increase in molecules between time points divided by the time interval, followed by the calculation of the average.

### Droplet-based measurements - Assessment of affinity

The binding strength of the antigen-antibody interaction was indirectly assessed by adding the antigen to the IgG bioassay, as previously described [48, 49]. In short, if the secreted antibody was specific for the antigen, the antigen relocated onto the nanoparticles. To assess this interaction, the fluorescence relocation of the IgG probe (≥LoD and ≤cutoff, as defined above) and the TT-APC relocation were plotted against each other, and the resulting slope of the curve was used as an indicator for affinity. To be defined as an antigen-specific cell, the fluorescence relocation of the antigen had to be above the LoD for at least one-time point:

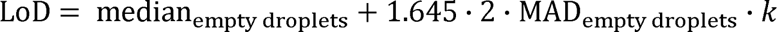

whereby median_empty droplets_ was the median relocation in droplets without a cell, MAD_empty droplets_ the median absolute deviation in droplets without a cell and *k* = 1.4826. The LoD was calculated for each experiment individually.

### Droplet-based measurements - Assessment of ROS production

To determine the ROS production level of the cells, the increase in their cell signal due to ROS-mediated oxidation and subsequent DNA binding of the reagent was measured over time. The increase in the signal served as an indicator of the level of ROS production and was calculated until the cell reached its maximum cell signal. Cells were categorized into ROS^low^ and ROS^high^ cells based on the distribution of signal increases measured on day 14 as following (see SI section 6):

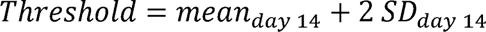

In specific experiments, *N*-acetylcysteine (NAC, 10 mM, Thermo Fisher, C10492) was added to the CellTrace Violet staining solution as well as to the aqueous phase I (20 mM) to serve as a ROS-negative control.

### Droplet-based measurements - Assessment of intracellular pH

Intracellular pH was measured as cellular fluorescence using the pH-responsive dye as described above. To calibrate the cellular signal indicating the intracellular pH (pH_i_), primary B cells were stained with pHrodo Green AM and CellTrace Violet and their pH_i_ was adjusted using the intracellular pH calibration buffer kit (Thermo Fisher, P35379) according to the manufacturer’s protocol. The cell signals were normalized to the median signal at pH 7.5 for calibration (SIFigure 10). To determine the pH_i_ of primary murine B cells, the cell signal was measured at the first time point. Here, the signal was normalized to the median cell signal of all encapsulated cells (assumed around physiological pH_i_. For the results, the cells were binned into two categories, acidic (pH_i_ <threshold) and physiological pH_i_, based on the calibration as follows:

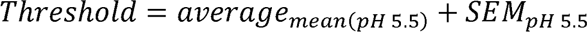

### Droplet-based measurements - Assessment of caspase-3/7 activity

The employed caspase substrate resulted in a cellular fluorescence signal increase when active enzymes were present in the droplet. The SD of the whole droplet fluorescence signal (SD_WDF_) was used to assess whether caspase-3 and 7 were activated (SIFigure 11). Based on the SD_WDF_ of droplets without a cell measured on the first time point, a threshold for detecting activated caspase-3/7 was defined as follows:

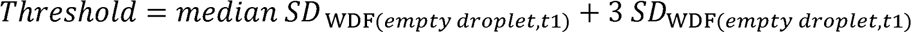

The threshold was calculated for each experiment individually.

### ELISpot

The number of B cells secreting IgG was quantified using a mouse ELISpot kit (Mabtech, 3825-2H). The isolated cells were counted using the Countess Automated Cell Counter II (ThermoFisher) and re-suspended at 0.5 x 10^6^ cells/mL in RPMI 1640 (Gibco, 11835-105) supplemented with 10% heat-inactivated FBS (Gibco, 10270-106), 1X Penicillin-Streptomycin (Gibco, 15140-122) and 25 mM HEPES (Gibco, 15630-080). ELISpot PVDF plates (Merck, MSIPS4W10) were activated using 15 μL/well of 35% ethanol, washed five times with sterile bi-distilled water (PanReac, A4042), and coated overnight with 15 μg/mL anti-IgG at 4 °C. Plates were washed five times with sterile PBS (Gibco, 10010-015) and 100 μL/well of the cell suspension was added and incubated at 37 °C for 24 h. The following day, the plates were emptied and washed five times with PBS. Total IgG was detected using 100 μL/well of anti-IgG-biotin antibody diluted at 1 μg/mL in PBS with 0.5% FBS and filtered through a 0.22 μm syringe filter (TPP, 99722) and incubated for 2 h at room temperature. Plates were then washed as previously described, and 100 μL/well of Streptavidin-HRP diluted at 1:1000 in PBS with 0.5% FBS was added for 1 h at room temperature. After washing the plates, 100 μL/well of TMB (Merck, T0565) was added to develop until distinct spots emerged. The reaction was stopped by washing extensively with deionized water, leaving the plates to dry. The plates were analyzed with AID ELISpot software 7.0 (Autoimmun Diagnostika).

### scRNA-Seq

#### scRNA-Seq - Sample and library preparation for mRNA sequencing

Cells isolated from the bone marrow and spleen of two mice were pooled on days 0, 3, 7, and 14 after secondary immunization. Cell preparation for single-cell sequencing was first performed, which included cell count, washing and re-suspension in 1x PBS with 0.04% BSA. Next, 16’000 single cells per sample were used to generate the 10x libraries using the Chromium Single Cell 5’ Reagent Kits v2 (Dual Index) (10x Genomics), following the manufacturer’s recommendations. After library preparation, all samples were sequenced on the NextSeq 2000 on a P2 100 cycles flow cell (136 bp paired-end).

#### scRNA-Seq - Pre-processing and processing of scRNA-Seq data

The SUSHI data analysis tool developed by the Functional Genomics Center (ETH Zurich/ University of Zurich) was used to process the sequenced samples [91]. The CellRanger application (v7.1), available on SUSHI, was used in the different processing steps. In short, de-multiplexing, unique molecular identifier (UMI) counting, and alignment to the mm10 mouse transcriptome were performed. Raw data generated by CellRanger were then read into the Seurat R package [92], also available on SUSHI. Next, the data were subjected to quality control using the Fastqc10xApp and FastqScreen10xApp, and normalization with the CellRangerApp on SUSHI. Thereby, parameters such as the total number of counts and the proportion of counts in mitochondrial genes were checked, and the removal of poor-quality cells (e.g., loss of cytoplasmic RNA). Furthermore, data normalization included the log-normalization of gene expression.

Subsequently, the normalized data were analyzed on the Loupe Browser application [93], which provided an interactive visualization functionality and allowed to identify and gain insights into the gene expression information of different cell populations/ clusters. These clusters were generated by using the t-distributed stochastic neighbor embedding (t-SNE) method [94] and the gene-based criteria are detailed in SITable 1. SIFigure 1 shows the population distribution, and SITable 2 shows their frequency over time. Furthermore, we identified IgG- and IgM-expressing cells (IgG- and IgM-ECs, respectively), i.e., plasma cells (PCs) that expressed *IGHG* (all isotypes) and *IGHM*, respectively. Their frequencies are detailed in SITable 3.

Differentially expressed genes (DEGs) for each cluster were identified using the Wilcox test in Seurat (p-value <0.05). The log2 fold changes (log2 FC) were used to compare the expression levels of each gene between the clusters and, over time, in the spleen and bone marrow separately. Shortly, the log2 FC value is the ratio of the normalized mean gene counts in each cluster relative to all other clusters for comparison, i.e., if log2 FC is 2 for a gene in cluster A *vs*. cluster B, the expression of that gene is increased in cluster A relative to B by a multiplicative factor of 2^2^ = 4.

#### scRNA-Seq - Data analysis

The list of DEGs obtained for each cluster and time point was used as input for pathway over-representation analysis (ORA) using the Consensus Path Data Base (CPDB) release 35 [95], considering a cutoff of 0.05. The Reactome database version 84 [96] and Kyoto Encyclopedia of Genes and Genomes (KEGG) [97] were selected as preferred databases for pathway analysis. The most relevant metabolic pathways were identified, as well as the DEGs (p-value <0.05) involved in those pathways and their respective expression level (based on log2 FC). A brief explanation of how these metabolic pathways, particularly glycolysis, lactate metabolism, and TCA cycle, were characterized is available in SI and SIFigure 2. A similar approach was used to investigate the activity of apoptosis and oxidative stress in B cells.

Regarding the activity of the metabolic pathways, they were investigated in IgG^low^- and IgG^high^-expressing cells (ECs) and IgM^low^- and IgM^high^-ECs (based on *IGHG* and *IGHM* expression levels), compared to all pooled B cells over time. Categorizing was performed based on low (log2-fold change (FC) ≤1) and high expression levels of IgG (log2 FC >1) as well as low and high levels of IgM (same threshold applied). The threshold was calculated based on the mean of log2 FC of *IGHG* and *IGHM*. The average log2 FC of the DEGs involved in the relevant metabolic pathways was considered to estimate their activity.

Next, a correlation between lactate metabolism, apoptosis and oxidative stress was investigated in low and high lactate-generating splenic B cells over time (based on *LDHA* expression levels, see SI section 1). Here, the activity of apoptosis and oxidative stress were also estimated based on the average log2 FC of the DEGs involved in the pathways. All heat maps and graphs were generated using R [98] and Microsoft Excel.

### Flow cytometry experiments

After the cell isolation, cells were re-suspended in PBS and treated with anti-CD16/CD32 (Clone 93, Biolegend, 101301, 1:200) for 15 min on ice. In parallel, live/dead staining was done using Zombie-Aqua (Biolegend, 423101, 1:500). Samples were stained in FACS buffer (PBS containing 2% FCS (Thermo Fisher) and 2 mM EDTA (Sigma-Aldrich)) for further 20 min on ice with the following antibodies: rat anti-mouse CD19 PE (Biolegend, 115507), rat anti-mouse CD138 FITC (Thermo Fisher, MA5-23554), rat anti-mouse Ly-6G Brilliant Violet 650 (Biolegend, 127641), rat anti-mouse Ly-6C APC/Cyanine7 (Biolegend, 128026), rat anti-mouse F4/80 APC (Biolegend, 123116), rat anti-mouse CD90.2 PerCP (Biolegend, 105321), hamster anti-mouse CD11c APC (Biolegend, 117310), rat anti-mouse I-A/I-E Brilliant Violet 421 (Biolegend, 107632) and rat anti-mouse CD45 Alexa Fluor 700 (Biolegend, 103128). Samples were acquired on a CytoFLEX flow cytometer (Beckman Coulter) using CytExpert software and the data were analyzed using FlowJo software 10.8.1.

