## Supporting Information for "Single-B cell analysis correlates high-lactate secretion with stress and increased apoptosis"

### **1. Characterization of the functionality bioassay to quantify the secretion of antibodies and to assess their interaction strength with antigen**

The bioassay to quantify IgG secretion rates of individual B cells and to assess the affinity of the secreted IgG toward the antigen has previously been published by our group [48, 49]. Here, the assay was adapted to our model system. For the detection of IgG, an alternative probe exhibiting similar binding strength for all IgG subtypes and none for TT was used (SouthernBiotech, 1033‑30, labeled with Alexa Fluor 488). To measure IgM secretion, an IgM probe was used (Jackson ImmunoResearch, 715-476-020, labeled with DyLight 405). The antigen used for immunizations, recombinant tetanus toxin heavy chain fragment C (FinaBio), was in-house labeled with APC (APC Conjugation Kit – Lightning-Link®, Abcam, ab201807; according to manufacturer's protocol).

SIFigures 1A and B show the fluorescence relocation of the anti-IgG and -IgM probes as a function of IgG and IgM antibody concentration, respectively. The limit of detection (LoD), cut-off, and plateau were calculated as described in the methods. The resulting analytical ranges were 3-375 IgG/s (0.3-15.0 nM, with 6 measuring points with 10 minutes intervals each), and 7-150 IgM/s (0.7-6.1 nM, with 6 measuring points with 10 minutes intervals each), respectively. B cells with a secretion rate ≥3 IgG/s were defined as IgG-secreting cells (IgG-SCs), and cells with secretion rates ≥7 IgM/s as IgM-SCs.

The fluorescence relocation of TT as a function of the concentration of a commercial anti-TT-specific IgG (Kerafast, EFD006) is shown in SIFigure 1C. A slope is obtained by plotting the fluorescence relocation of TT against the fluorescence relocation of the IgG probe (above LoD and below cut-off, SIFigure 1D). Previously, our group showed that this slope correlated with affinity and, more specifically, the dissociation constant of the interaction [42]. In short, the slope indicated the interaction strength between the secreted IgG and antigen, and increased affinity resulted in increased slopes. Here, a slope of 1.64 was obtained for the commercial TT-specific IgG.

| A | B |
| --- | --- |
| 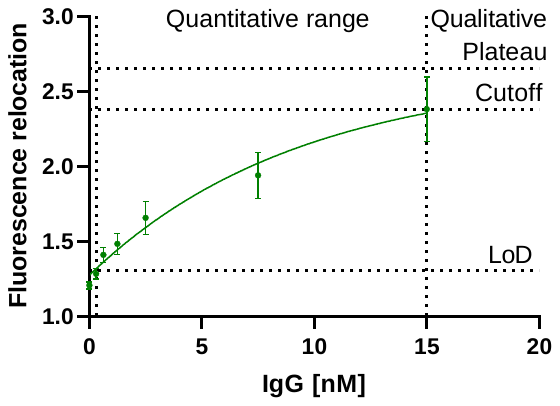 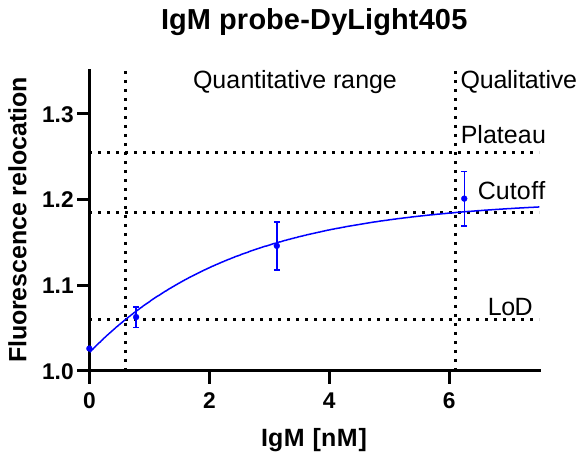 | |
| C | **D** |
| 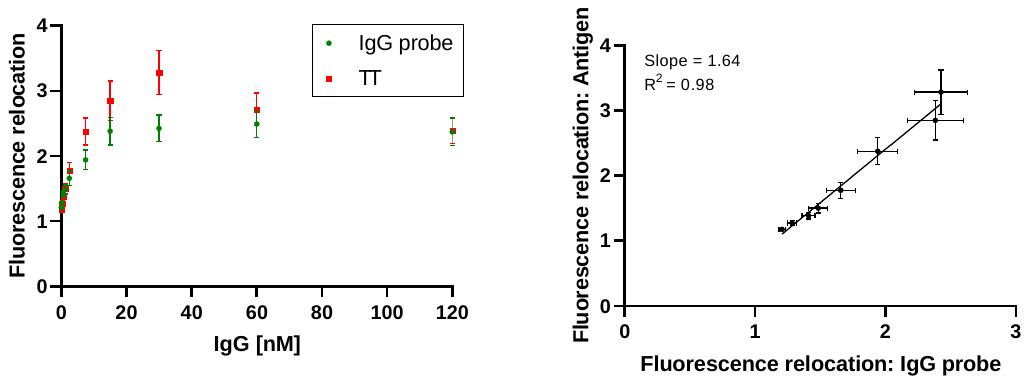 | |

SIFigure 1: (a) Fluorescence relocation of the goat anti-mouse IgG antibody labeled with Alexa Fluor 488 as a function of the IgG concentration. When measuring six time points with an interval of 10 min, the resulting quantitative range is from 3-375 IgG/s. Cells with a secretion rate ≥3 IgG/s are defined as IgG-secreting cells. The LoD, the cut-off and the plateau are shown. (b) The change in fluorescence relocation of the donkey anti-mouse IgM F(ab')2 labeled with DyLight 405 concerning the IgM concentration. By measuring six time points with a 10-minute interval, a quantitative range of 7-150 IgG/s was obtained. Cells that secrete IgM at a rate ≥7 IgM/s are classified as IgM-secreting cells. The LoD, cut-off, and plateau are depicted. (c) The relationship between the fluorescence relocation of the IgG probe and the APC-labeled antigen, TT, and the concentration of a TT-specific IgG. (d) Fluorescence relocation of TT as a function of fluorescence relocation of the IgG probe. A slope representing the interaction strength is obtained by plotting the fluorescence relocalizations of the antigen and IgG probe against each other. The slope of the TT-specific antibody used here is 1.64.

### **2. Further elaboration on transcriptomic analysis**

SITable 1: Key surface and intracellular gene markers that were applied to identify subpopulations of interest in spleen and bone marrow.

| Plasma cells | *SDC1* (CD138), *CXCR4, PRDM1* (BLIMP1), *XBP1* |
| --- | --- |
| IgG-ECs | Plasma cells expressing *IGHG* |
| IgM-ECs | Plasma cells expressing *IGHM* |
| Activated B cells | *CD19, MS4A1* (CD20), *CD27, CD80, PAX5* |
| GC B cells | *CD19, CD37,* *MS4A1* (CD20), *CCR6, BCL6* |
| Memory B cells | *CD19, CR2* (CD21), *NT5E* (CD73), *POU2AF1* (OBF1), *SPI-B* |
| Immature B cells | *CD19, CD24, CD93, PTPRC* (B220), *PAX5, EBF1* |

SITable 2: Frequency of the different cell populations in the spleen on days 0, 3, 7 and 14 as identified through single-cell transcriptomic analysis, and whose clusters are represented in SIFigure 2. The scRNA-Seq data was generated from cells of two mice pooled in a ratio of 1:1 prior to sequencing.

|  | Day after secondary immunization | | | |
| --- | --- | --- | --- | --- |
|  | **0** | **3** | **7** | **14** |
| Plasma cells | 9% | 26% | 31% | 33% |
| Activated B cells | 14% | 9% | 12% | 10% |
| GC B cells | 36% | 27% | 19% | 35% |
| Memory B cells | 9% | 21% | 25% | 22% |
| Immature B cells | 30% | 17% | 13% | 0% |
| Others | 2% | 0% | 0% | 0% |

| A  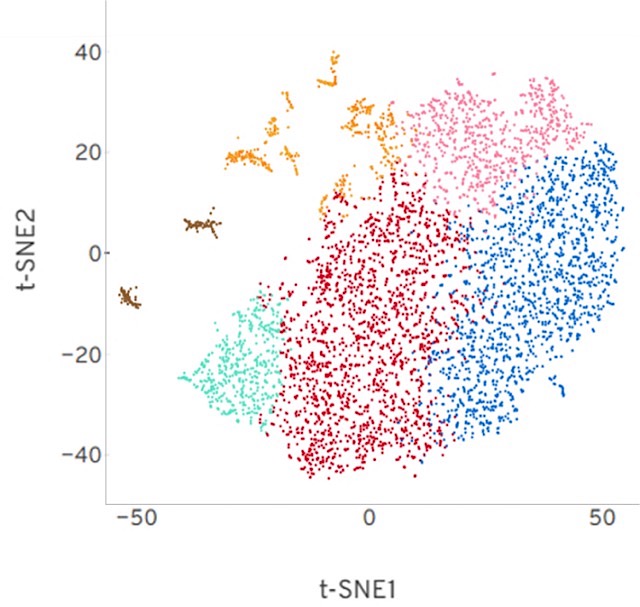 | B  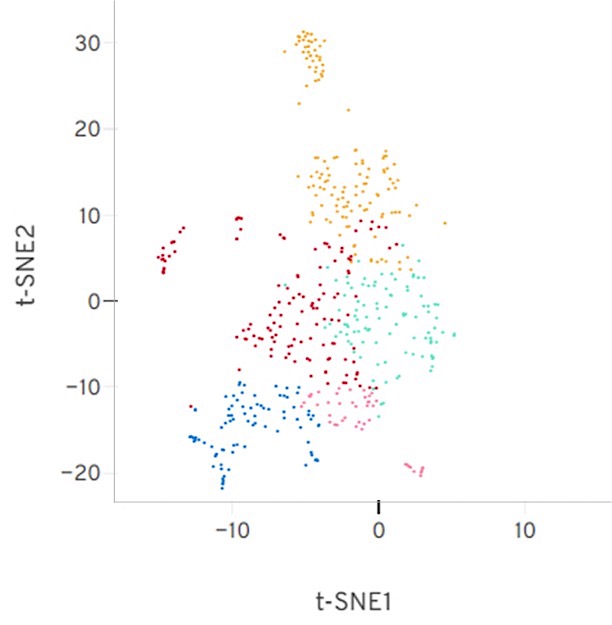 |
| --- | --- |
| C  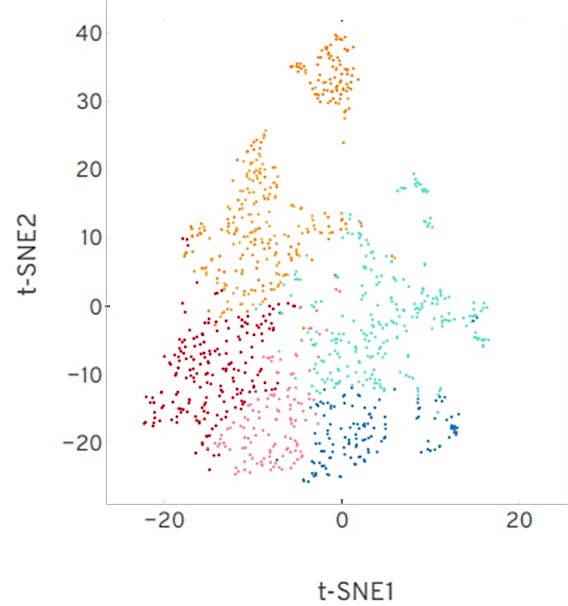 | **D**  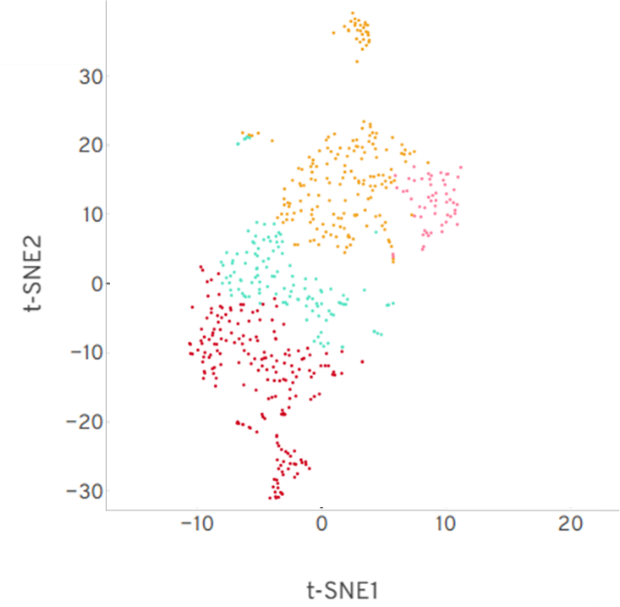 |

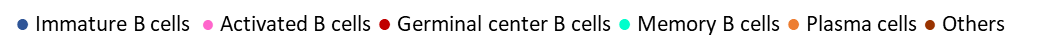

SIFigure 2: Identification and t-SNE projections (Loupe Browser) of the different cell populations in the spleen on (a) day 0, (b) day 3, (c) day 7 and (d) day 14. Legend: Immature B cells in blue; Activated B cells in pink; GC B cells in red; Memory B cells in light green; Plasma cells in orange; Other cells (i.e., non-B cells) in brown.

SITable 3: Frequency of IgG- and IgM-ECs in the spleen and bone marrow throughout the immune response obtained by transcriptomic analysis. The scRNA-Seq data was generated from cells of two mice pooled in a ratio of 1:1 prior to sequencing.

|  |  | Day after secondary immunization | | | |
| --- | --- | --- | --- | --- | --- |
|  |  | **0** | **3** | **7** | **14** |
| Spleen | **IgG-ECs** | 0.3% | 1.4% | 0.7% | 0.9% |
|  | **IgM-ECs** | 0.8% | 3.0% | 3.9% | 4.0% |
| Bone marrow | **IgG-ECs** | 0.1% | 0.1% | 0.3% | 0.3% |
|  | **IgM-ECs** | 1.1% | 1.7% | 1.8% | 1.6% |

All splenic B cells were also classified based on their *LDHA* expression levels so that we could study low and high lactate-generating cells. B cells with a log2 FC ≤0 were classified as low lactate-generating cells and >0 as high lactate-generating cells.

Regarding the expression analysis of genes involved in glycolysis, lactate metabolism (generation) and TCA cycle, SIFigure 3 illustrates how these pathways were defined for the transcriptomic analysis. In short, glycolysis starts from glucose being metabolized into glucose-6-phosphate and continues until the generation of pyruvate. Lactate metabolism considers the generation of lactate from pyruvate. Lastly, the TCA cycle starts with the formation of acetyl-CoA. By doing so, we could distinguish whether B cells preferred the pathway that leads to lactate or to fuel the TCA cycle. The rest of the metabolic pathways, namely pentose phosphate pathway (PPP), oxidative phosphorylation (OXPHOS), fatty acid (FA) synthesis and β-oxidation (FAO), and glutaminolysis, as well as apoptosis and oxidative stress, were based on the Kyoto Encyclopedia of Genes and Genomes (KEGG) database.

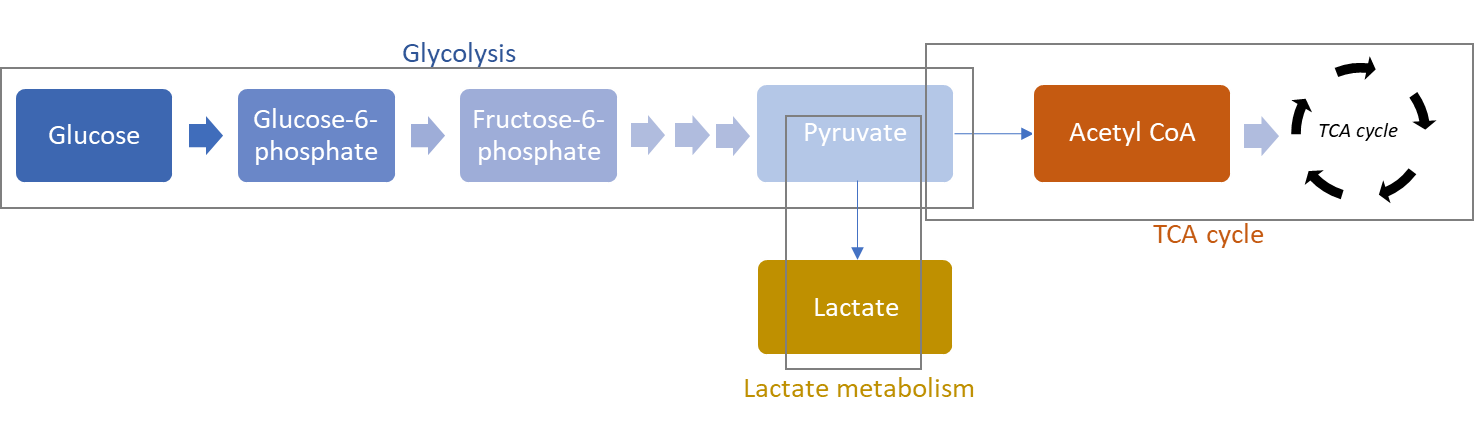

SIFigure 3: Illustration of how glycolysis, lactate metabolism and TCA cycle were considered as metabolic pathways for gene expression analysis.

### **3. Characterization of the lactate assay**

A commercial fluorometric assay kit (Lactate Assay Kit, Sigma-Aldrich, MAK064) was used for the lactate measurement. The enzyme concentration was adapted to measure the secreted lactate in real-time. For this purpose, droplets with defined lactate concentrations were prepared, and the fluorescence increase was measured over time to estimate the kinetic of the reaction. Using an increased enzyme concentration (4x of manufacturer's concentration, i.e., 8 μL enzyme mix per 100 μL bead solution), the highest lactate concentration used (200'000 amol/nL) led to the full conversion of the fluorescent substrate in less than 10 min. Since the lactate secretion rate of B cells was estimated to be much lower, this enzyme concentration appeared suitable to ensure the full conversion of the secreted lactate measuring of single B cells.

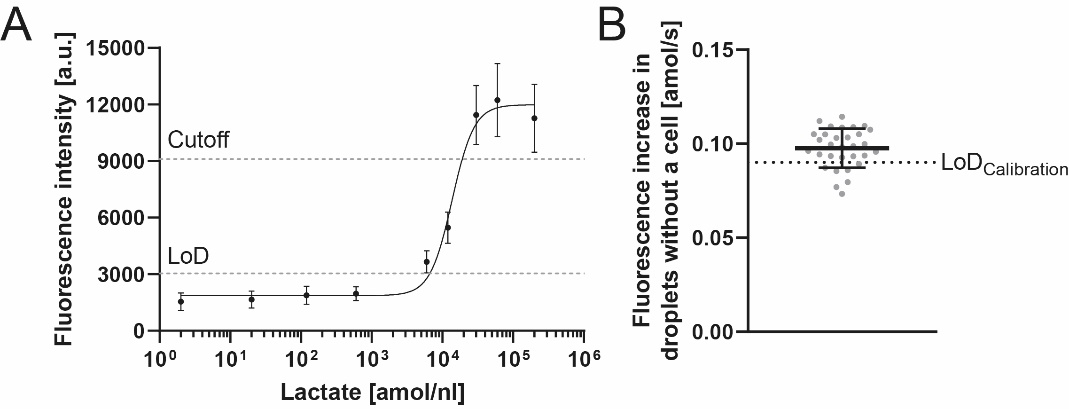

SIFigure 4: (a) Fluorescence intensity of the droplet as a function of the lactate concentration. The analytical range of concentrations corresponds to 0.09-0.80 amol/s when measuring 6 times with an interval of 10 min. (b) The detection limit (LoD) of the lactate assay defined by the calibration (0.09 amol/s, dashed line) was adjusted due to a background level of lactate accumulating during the encapsulation process by secreting cells, together with further limitations due to photochemistry and diffusion of lactate and the fluorescent product. The points represent the average of the droplets without a cell of individual measurements of spleen and bone marrow samples performed throughout the immune response (n= 32). The average fluorescence increase corresponded to a secretion rate of 0.10 ± 0.01 amol/s. Accordingly, the quantitative range of the lactate bioassay was adjusted to 0.10-0.80 amol/s.

A limitation of the detection limit of the bioassay was observed in a control experiment using primary B cells. Here, the fluorescent signal in droplets not containing a cell increased over time, corresponding to a median secretion rate of 0.10 amol/s. The following three causes could be identified for the increase in fluorescent signal, i.e., photochemical activation and diffusion of lactate and fluorescent product. When combining assays and defining the measurement time, these limiting factors should be considered. To observe potential measurement issues, we always used the droplets not containing a cell as an internal control. The average increase in the empty droplets for all included measurements was 0.10 ± 0.01 amol/s (SIFigure 4, mean ± SEM) and thus resulted in only a small increase of the detection limit compared to the one defined based on the calibration.

*Photochemical activation of the fluorescent probe*

First, the selected light intensities and exposure times for the excitation of fluorophores were shown to lead to a photochemical reaction, increasing the fluorescence signal of the lactate assay. The influence of photochemical activation on the determination of secretion rates could be reduced by decreasing the concentration of the fluorescent probe (0.5x of manufacturer's concentration, i.e., 0.5 μL fluorescent probe per 100 μL bead solution). By optimizing the concentration of the probe and the settings of the light source, we could decrease the observed photoactivation over six measurement points corresponding only to a 'secretion rate' of 0.06 ± 0.01 amol/s. Thus, the photoactivation and its influence on measurement were below the LoD, which was defined based on a calibration with lactate standard.

*Diffusion of the fluorescent product*

Next, we controlled whether the fluorescent product diffused from one droplet to another. For this purpose, droplets containing the lactate assay and 10'000 amol/droplet lactate were prepared and incubated for 10 min to ensure complete conversion. Afterward, the droplets were mixed with droplets lacking the lactate assay, introduced into the chamber and measured. The 'empty' droplets contained calcein, which was previously shown to not diffuse between droplets (internal lab data), and its green fluorescence was used for identification. After mixing the two distinct droplet populations, the mixture was introduced into an observation chamber, and the calcein-positive droplets' fluorescence intensity for the fluorescent product of the assay was measured over time. Indeed, these droplets slightly increased in fluorescence over time, indicating diffusion of the fluorescent product. The increase corresponded to a median secretion rate of 0.15 ± 0.08 amol/s.

*Diffusion of lactate*

Diffusion of lactate was demonstrated using a mixture of droplets containing the lactate kit and calcein mixed with another distinct droplet population containing a high lactate concentration (600'000 amol/nL). Although the droplets containing the assay did not contain lactate, an increase in fluorescence corresponding to a secretion rate of 0.27 ± 0.07 amol/s was observed. However, we assumed this factor to be less important for cell measurements as no such high concentration of secreted lactate was measured using B cells, and the lactate would be converted quickly by the present assay components in cell measurement. Nonetheless, lactate might diffuse between droplets if not efficiently converted quickly.

### **4. Additional results from the cell measurements**

| A | B | C |
| --- | --- | --- |
| 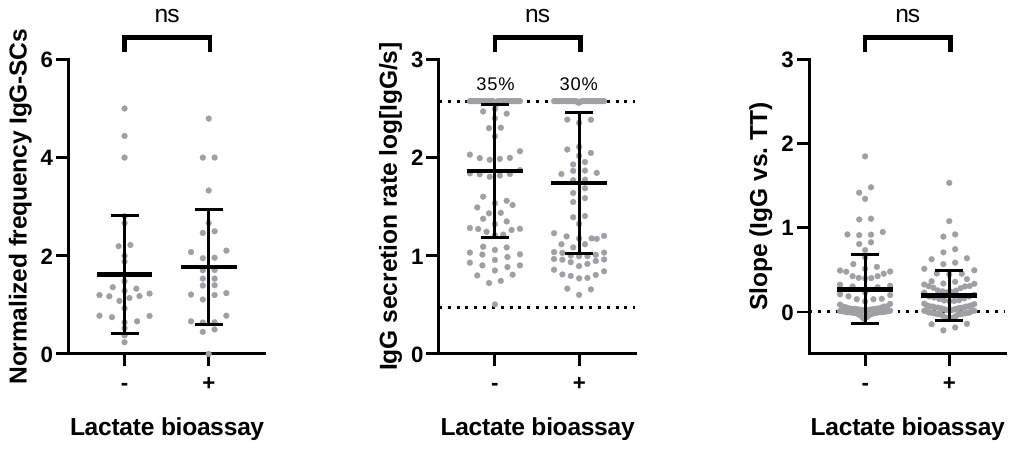 | | |

SIFigure 5: Combination of the functionality and lactate bioassays did not lead to significant deviations of (a) the frequency of IgG-secreting cells, (b) distributions of IgG secretion rates and (c) slopes, the indicator of the interaction strength between the secreted IgG and antigen. For (a), the IgG-SC frequencies were normalized by the respective frequency measured using ELISpot. For (b) and (c), a randomly-selected uniform number of data points were pooled from measurements of spleen and bone marrow samples from 4 mice per day. The number above the dotted line represents the frequency of cells with a secretion rate equal or above the assay’s quantitative range. P values were 0.33, 0.17 and 0.50, respectively.

| A  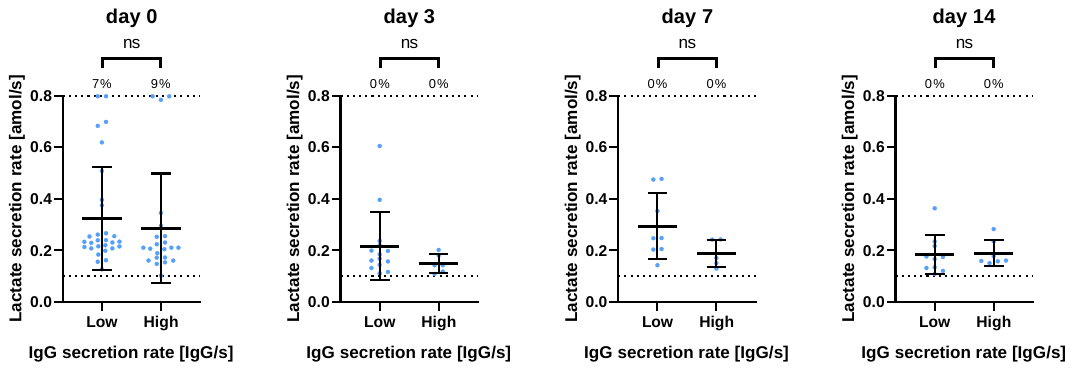 | |
| --- | --- |
| B | **C** |
| 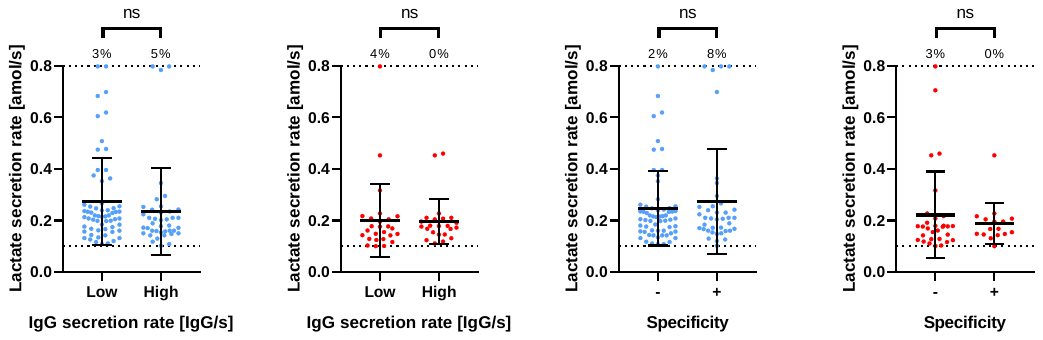 | |

SIFigure 6: (a) Comparison of the distribution of lactate secretion rates between IgG-SCs with high or low IgG secretion rates (threshold 200 IgG/s) on individual days in the spleen (blue). (b) Comparison of the distribution of LSRs between IgG^low^- and IgG^high^-SCs in the spleen (blue, left) and bone marrow (red, right). Data from the different measurement days were pooled. (c) Classification of IgG-SCs based on the slope into specific and non-specific cells. Comparison of the LSR distribution within the spleen (blue, left) and bone marrow (red, right). All distributions were non-significantly different. Data from four mice per time point were pooled. The number above the dotted line represents the frequency of cells with a LSR ≥0.8 amol/s.

### **5. Purity of splenic and bone marrow samples after cell isolation**

To obtain the B cells from the splenic and bone marrow samples, a depletion method using the Pan B Cell Isolation Kit II (Miltenyi Biotec) was employed. Flow cytometry was used to evaluate the frequency of B cells in the purified samples. In short, the B cells were identified based on the expression of CD19 or CD138, i.e., CD19^+^ or CD19^-^CD138^+^ as markers for B cells and plasma cells. The frequency of B cells in the purified samples was 95 ± 2% in the splenic samples and 49 ± 27% in the bone marrow samples (n= 7, SIFigure 8A). Variations of bone marrow purities were distributed among analyzed days (days 0, 3 and 7). Variations in purity in bone marrow might alter the extracted frequencies of IgG-SCs (Figure 1B), but none of the other extracted parameters were influenced. Accordingly, the kit was efficient in isolating B cells from spleen samples but not from the bone marrow, and B cells from bone marrow were only studied functionally as IgG-SCs.

A refined division of the non-B cells is visible in SIFigure 8B (average of 5-7 measurements). The majority of non-B cells in the spleen samples were identified as CD45^+^ leukocytes that were not positive for the selected markers characteristic for non-B cell lymphocytes (see methods), followed by monocytes (Ly6C^+^Ly6G^-^), dendritic cells (MHCII^+^CD11c^+^) and T cells (CD90.2). In the bone marrow, the majority of non-B cells were CD45^+^ leukocytes that were not positive for the selected markers characteristic for non-B cell lymphocytes (see methods), followed by monocytes (Ly6C^+^Ly6G^-^) and neutrophils (Ly6C^+^Ly6G^+^). FMO controls were used for the definition of the gates (see SIFigure 8C for gating strategy).

In conclusion, phenotypic studies of the purified cells corresponded well with cells from the B cell lineage in the spleen. However, the phenotypic studies of bone marrow B cells are limited by the presence of other cells. Therefore, the analyses were only performed focused on subpopulations like ASCs, which are identified by the detection of antibody secretion, or the transcriptomic data where B cells were identified. Finally, the determined frequency of IgG-SCs relative to all encapsulated cells was influenced by the variable purity in the bone marrow.

| A  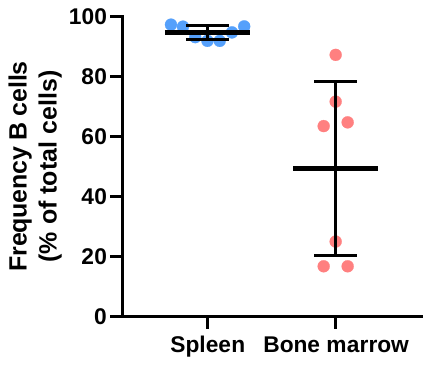 | B  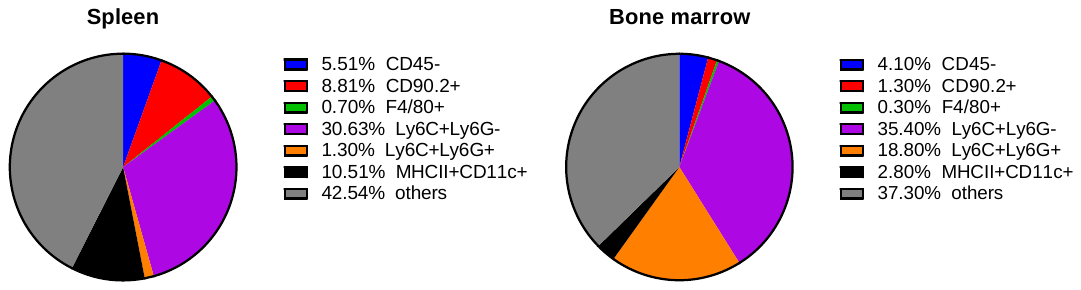 |
| --- | --- |
| C  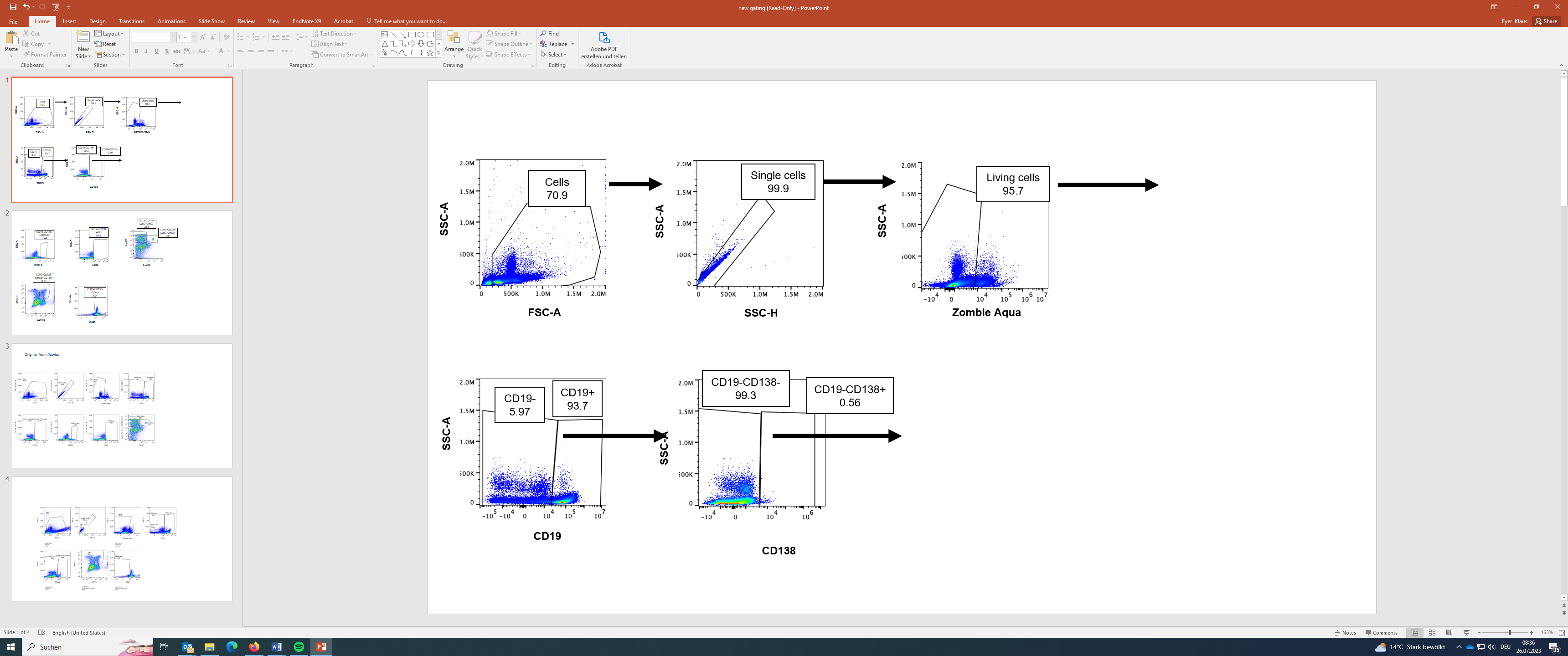 | |
| 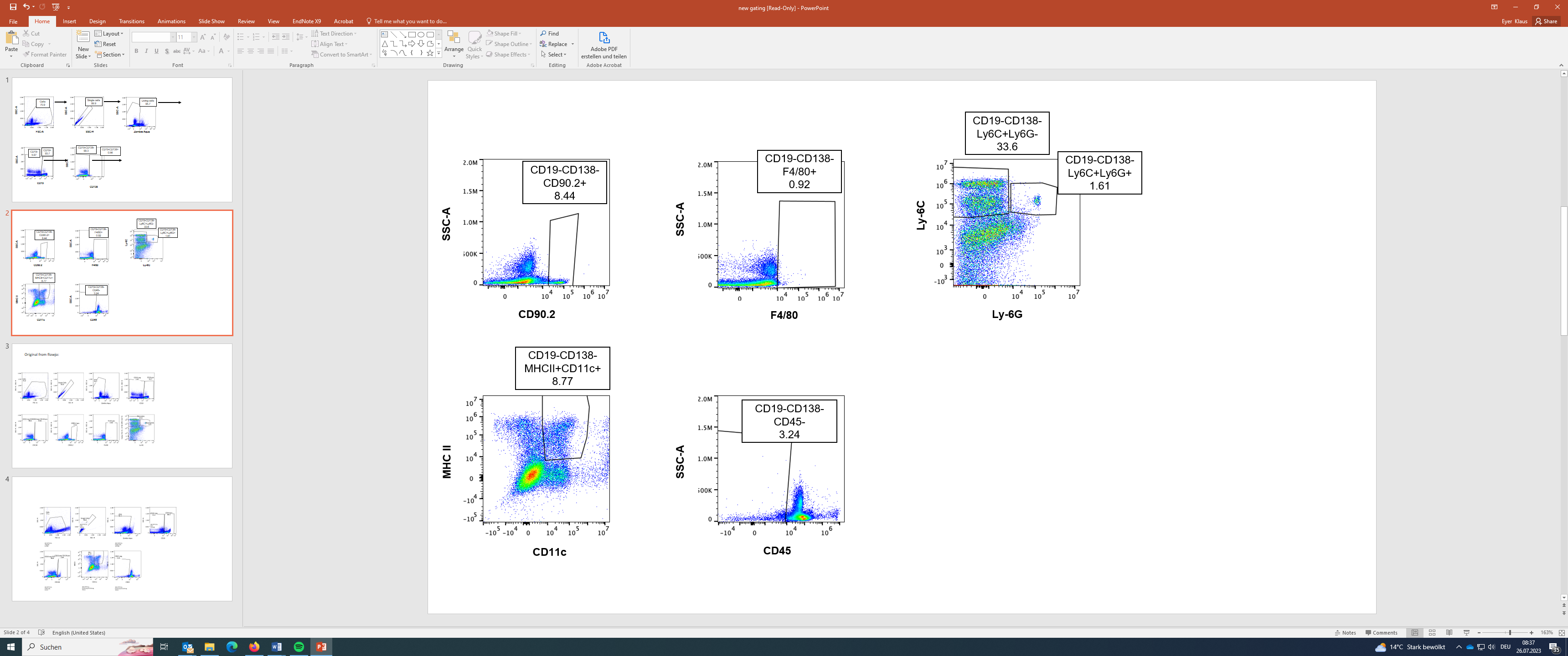 | |

SIFigure 7: Flow cytometry measurement of purified spleen and bone marrow samples. (a) Analysis of B cell purity in isolations from spleen and bone marrow. The frequency of cells from the B cell lineage in the purified samples was determined based on the expression of CD19^+^ and CD19^-^CD138^+^ (n= 7). (b) Distribution of non-B cells according to markers used (average of 5-7 measurements). (c) Gating strategy for the analysis of splenic non-B cells.

### **6. Comparison of frequency of splenic IgG-SCs**

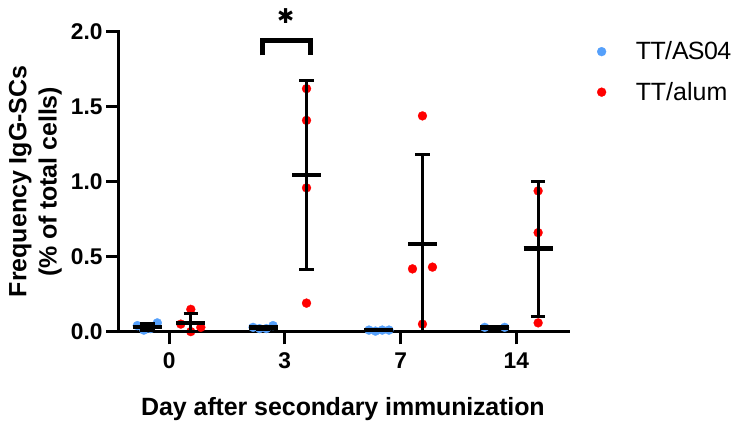

SIFigure 8: Frequency of splenic IgG-SCs. Immunization with TT/alum (i.e., without MPLA) resulted in significantly more IgG-SCs on day 3 compared to TT/AS04, and a tendency toward increased frequency was observed on days 7 and 14.

### **7. Characterization of the ROS bioassay**

The ROS level as an indicator of oxidative stress was measured by adding a cell-permeable, ROS-sensitive reagent (CellROX Green Reagent, Thermo Fisher, C10492) to the aqueous phase II during droplet production. The fluorescence intensity of the reagent changed from low to high intensity upon oxidation and subsequent binding to DNA resulting in a prominent signal (SIFigure 9A). The average linear fluorescence increase over time [a.u./10 min] was calculated to indicate ROS production up to each cell maximum (SIFigure 9B). Higher averages corresponded to higher production of ROS.

To further control the specificity of the assay, cells were treated with the ROS scavenger *N*-acetylcysteine (NAC, 10 mM, 30 min on ice; Thermo Fisher, C10492) prior to encapsulation. NAC was added to the aqueous solution I of the droplets (20 mM). The distribution of increases of fluorescence intensity over time of the NAC-treated cells was statistically different to the one of untreated cells (SIFigure 9C). As expected, much lower fluorescence increases were measured due to the scavenging effect of NAC.

At least two populations of cells were clearly distinguishable – cells with a fast and strong increase in signal (called ROS^high^) and cells with only little signal increase over time (ROS^low^) (SIFigure 9B). A threshold was defined to separate the populations and compare parameters like lactate secretion rates between them. The control with NAC was not suitable for the definition as NAC had ROS scavenging properties, leading to signal increases lower than physiologically observed (comparison to untreated cells). Therefore, we looked at the distribution of the signal increases over the course of the immune response (SIFigure 9D). On day 14, a relatively homogenous population with low signal increases was observed. Therefore, the threshold was defined based on their distribution as follows:

$$Threshold=mean_{day 14}+2 SD_{day 14}$$

The resulting 1'810 a.u./10 min threshold is displayed as a vertical line in SIFigure 9D. Throughout the immune response, 47% (day 0), 33% (day 3), 32% (day 7) and 6% (day 14) of the splenic B cells were classified as ROS^high^.

| A  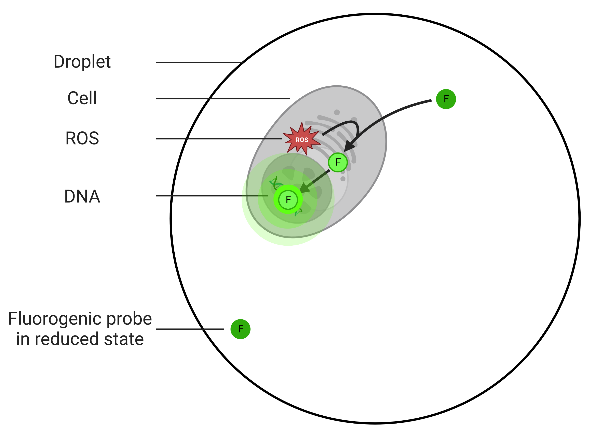 | B  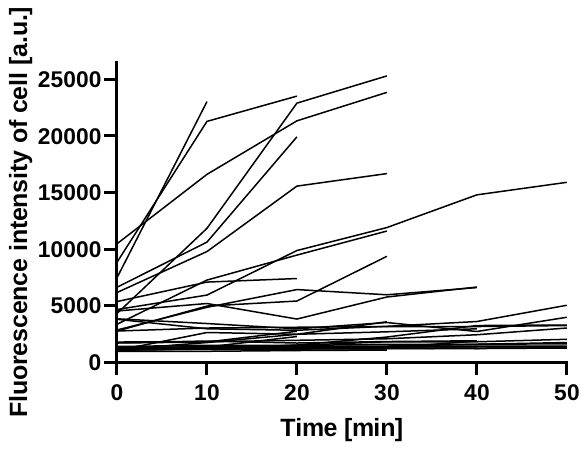 |
| --- | --- |
| C  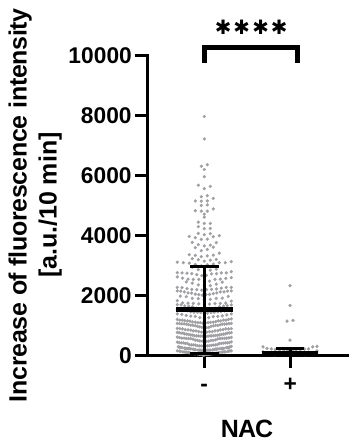 | **D**  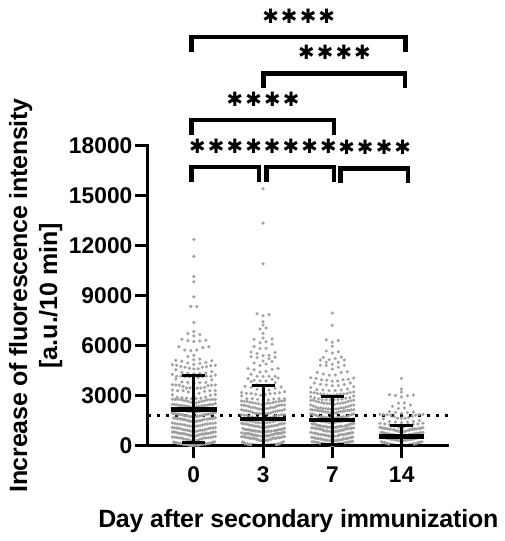 |

SIFigure 9: (a) Schematic representation of the bioassay used to assess cellular ROS production levels. A cell-permeable reagent is used, which increases its fluorescence intensity upon oxidation by ROS and binding to DNA. (b) Fluorescence signal of a selection of splenic B cells over time until the maximum cell signal is reached (n_traces_= 52). (c) Comparison of the distribution of ROS levels of untreated and NAC-treated splenic cells at day 7 (n= 375 per day, i.e., randomly selected 125 cells per mouse). (d) Distribution of the average increase in cell signal of splenic B cells over the time of the immune response. Each dot represents one individual cell. The vertical line represents the threshold used for categorizing the cells into ROS^low^ and ROS^high^ cells. The level of statistical significance is denoted as *p <0.05, **p <0.01, ***p <0.001 and ****p <0.0001. Panel A was created with BioRender.com.

### **8. Characterization of the bioassay indicating the intracellular pH**

Cells are stained with a pH-sensitive fluorogenic probe (pHrodo^TM^ Green AM, Thermo Fisher, P35373) to assess the intracellular pH (pH_i_). Thereby, the fluorescence intensity of the cell increased upon decreasing pH_i_ (SIFigure 10A). For calibration, we adjusted the pH_i_ to different extracellular pH values using a kit (intracellular pH calibration buffer kit, Thermo Fisher, P35379) and measured the fluorescence intensity of the cells (SIFigure 10B, mean and SEM of 4 individual measurements are shown). Indeed, the fluorescence intensity increased when pH was decreased, validating the dye as a pH_i_ indicator. The fluorescence intensity was measured at the first measuring time point. We decided to normalize the fluorescence intensities as we observed variability in the absolute fluorescence signals between calibration experiments (for example, due to different loading). Therefore, the cell signal measured at the first time point was normalized to the median of all cells, given that most cells clustered to one population (SIFigure 10C) and assumed that they have a physiological pH_i_.

In addition to correlation studies at the single-cell level, comparative studies of parameters such as LSR between cells with physiological and acidic pH_i_ were also performed. The calibration was used for the definition of the threshold. Due to the small differences between the normalized signals, we set the threshold as follows:

$Threshold=average_{mean\left( pH 5.5 \right)}+SEM_{pH 5.5}$ = 1.59

The frequency of cells with cell signal >1.59, i.e., acidic pH_i,_ was 6% on day 0, 4% on day 3, 4% on day 7, and 9% on day 14.

| A | B | C |
| --- | --- | --- |
| 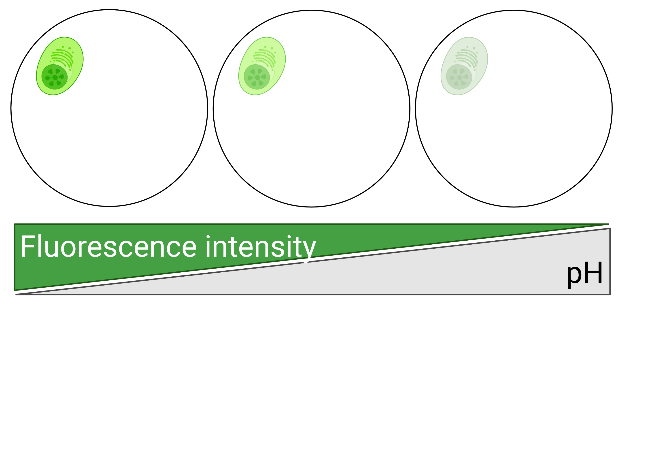 | 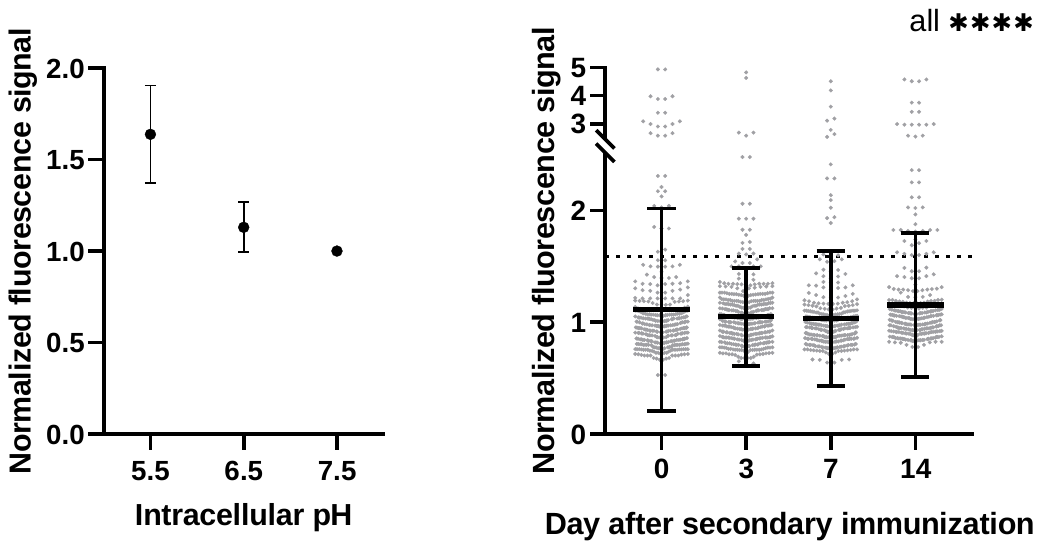 | |

SIFigure 10: (a) To assess the pH_i_, the cells were stained with a pH-sensitive dye. (b) The dependence of the fluorescence intensity on pH was demonstrated using primary B cells and different intracellular pH buffer solutions. The fluorescence intensities were normalized to the intensity measured at pH 7.5. (c) Distribution of normalized fluorescence signal of splenic cells throughout the immune response. The distributions differed significantly between all days (all p-value <0.0001). The vertical line represents the threshold. Panel A was created with BioRender.com.

### **9. Characterization of the bioassay to assess caspase-3/7 activity**

To assess caspase-3/7 activity, a fluorogenic substrate for these enzymes was used (CellEvent Caspase-3/7 Green Detection Reagent, Thermo Fisher, C10723). The substrate is intrinsically non-fluorescent and was added to the aqueous phase of the droplets. The probe was a four amino acid peptide (DEVD) with a cleavage site for caspase-3/7, which was conjugated to a nucleic acid-binding dye. Upon cleavage by activated caspase-3/7, the dye bound to DNA and resulted in a bright fluorescence (SIFigure 11A). The SD of the total droplet signal over time was used to identify cells with activated caspase-3/7 (termed caspase-3/7^+^ cells). SIFigure 11B displays traces of droplets containing a caspase-3/7^+^ cell (green, n= 5) as well as of traces of droplets with a caspase-3/7^-^ cells and without a cell (black, each n= 5). The threshold for classifying cells in cells with or without caspase-3/7 activity was defined for each measurement based on the droplets containing no cells as follows:

$$Threshold=median_{empty droplets(SD of droplet signal)}+3 SD_{empty droplets(SD of droplet signal)}$$

Application of this threshold resulted in <0.5% droplets classified as positive of total droplets not containing a cell.

| A  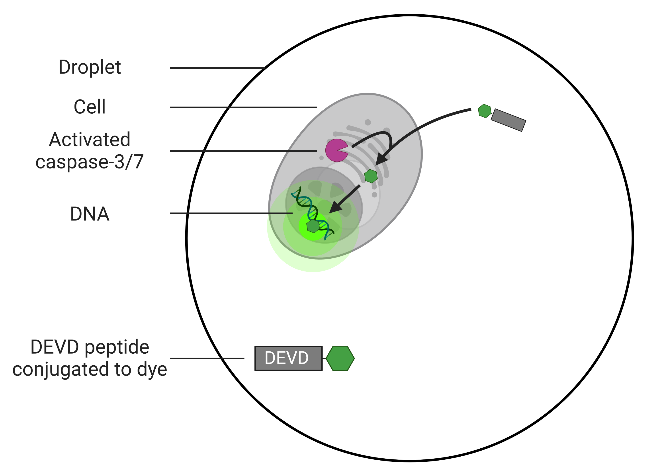 | | B  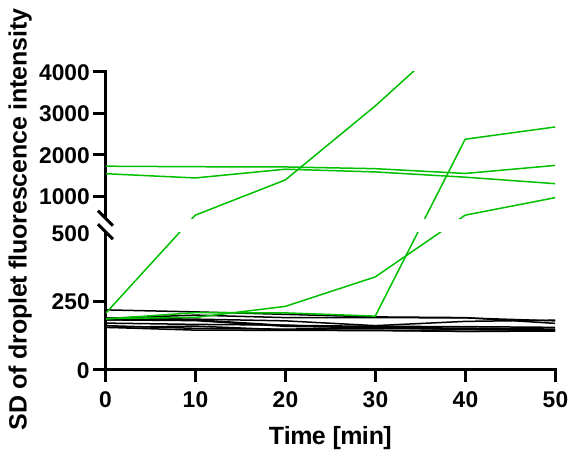 |
| --- | --- | --- |
| C | **D** | |
| 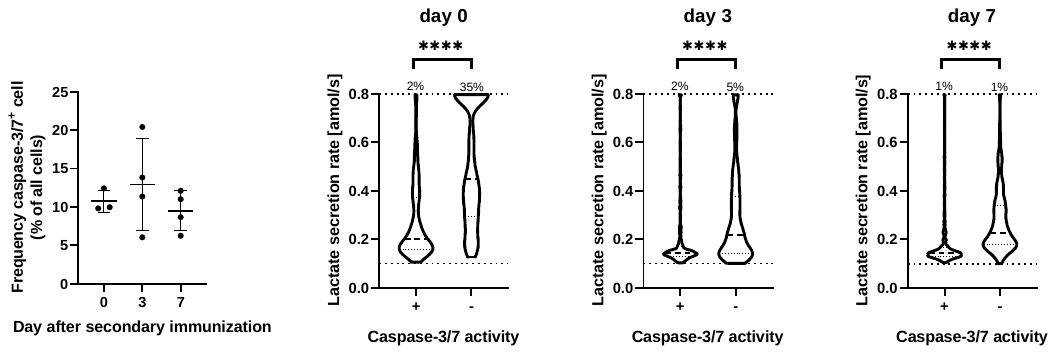 | | |

SIFigure 11: (a) Schematic representation of the bioassay for the detection of active caspase-3/7. Active caspase-3/7 cleaves the peptide-dye conjugate, allowing the free dye to bind to the DNA, resulting in a strong fluorescence. (b) Traces of caspase-3/7^+^ cells (green, n= 5) as well as droplets with cells negative for caspase activity (black, n= 5) and without cells (black, n= 5). Based on the droplets without cells, a threshold for the detection of active caspase-3/7 was defined. (c) Frequency of splenic cells displaying active caspase-3/7 on days 0, 3 and 7. The frequency did not vary significantly between the days. (d) The cells were categorized depending on observed caspase-3/7 activity and their distributions of LSRs compared. A random selection of 125 cells per mouse (3-4 mice) was pooled for every category. The percentage above the dotted line indicated the frequency of cells with a LSR ≥0.8 amol/s. Cells positive for caspase-3/7 activity showed a reduced median LSR and a greater prevalence of cells with a low LSRs (<0.2 amol/s, all p-values <0.0001). Panel A was created with BioRender.com.
